# Genome reshuffling as a route to specialization? Chromosome-level insights from the bark beetle symbiont genus *Geosmithia*

**DOI:** 10.64898/2026.09.15.751753

**Authors:** Tereza Veselská, Miroslav Kolařík, Ryan R Bracewell

## Abstract

Evolution of genome architecture is increasingly recognized as a major driver of fungal adaptation. However, because most studies have focused on plant and human pathogens, the genomic mechanisms underlying adaptation beyond pathogenic lifestyles remain poorly understood. Here, we establish the bark beetle–associated fungal genus *Geosmithia* as a model for studying genome evolution during ecological transitions. *Geosmithia* species occupy diverse ecological niches ranging from generalists, facultative symbionts of multiple beetle vectors and host tree species to more specialized conifer specialists and obligate nutritional symbionts of ambrosia beetles. Using chromosome-level genome assemblies of eleven species representing independently evolved ecological strategies, we investigated how genome architecture, repetitive DNA, and gene family evolution contribute to this ecological diversification. Comparative genomics revealed pervasive intra- and interchromosomal rearrangements, even between recently diverged species, demonstrating that extensive chromosome restructuring can accumulate over short evolutionary timescales. Breakpoints frequently occurred in gene-rich rather than repeat-rich regions, contrasting with the transposable element-associated patterns commonly described in fungi. Hi-C analyses revealed a typical Rabl chromosome organization but highly unusual centromeres that are repeat-poor, gene-rich, transcriptionally active, and conserved as syntenic blocks across species. Further, centromere identity may be determined by a highly divergent CenH3 variant. Despite similar genome sizes, specialists had increased repetitive DNA content and widespread gene family contractions, consistent with metabolic streamlining during ecological specialization. Our results demonstrate that extensive structural genome evolution can occur in gene-dense fungal genomes and establish *Geosmithia* as a powerful model for studying the interplay between genome evolution and ecological diversification.

## Introduction

Fungi are fundamental components of ecosystems, playing key roles in nutrient cycling, decomposition, and ecosystem productivity, while linking many organisms to one another through symbiotic interactions (Bahram and Netherway, 2022; Peay et al., 2016; Treseder and Lennon, 2015). Beyond their ecological importance, fungi also represent a major source of biotechnological innovations, including the production of enzymes, pharmaceuticals, bioactive compounds, and industrial biomaterials (see Niego et al., 2023). The ecological success of fungi largely stems from their remarkable evolutionary plasticity and capacity for evolutionary innovations (Mestre et al., 2020; Naranjo-Ortiz and Gabaldón, 2020; Sauters and Rokas, 2025). Comparative genomics has revealed that fungal adaptation can be driven not only by single nucleotide substitutions, but also by large-scale structural genome evolution and specific gene family expansions and contractions (Hartmann, 2022; Sauters and Rokas, 2025).

Recent comparative genomic analyses using chromosome-level genomes assemblies have shown that extensive structural variation can accumulate even between closely related fungal species (Ament-Velasquez et al., 2024; Shi-Kunne et al., 2018; Treindl et al., 2021) and may contribute to adaptation, reproductive isolation, and host specialization (Li et al., 2023; Rajeh et al., 2018). Chromosomal rearrangements in fungi have been associated with pathogenicity, stress tolerance, host adaptation, and niche specialization (Ament-Velasquez et al., 2024; Coelho et al., 2025; Quenu et al., 2022; Shi-Kunne et al., 2018; Todd et al., 2017; Treindl et al., 2021). In plant pathogenic fungi, rearrangements are frequently linked to repeat-rich genomic regions and accessory chromosomes that facilitate rapid evolution of virulence-associated genes (Ma et al., 2010; Schotanus et al., 2015; Seidl et al., 2020).

Large-scale structural variation, such as chromosome fusions and fissions, often require subsequent stabilization because they can generate dicentric or acentric chromosomes. In fungi, the evolutionary plasticity of centromeres, including centromere inactivation and neocentromere formation, provides mechanisms that stabilize these rearranged chromosomes and facilitate their long-term persistence (Dutta et al., 2025; Guin et al., 2020; Narayanan et al., 2024; Sankaranarayanan et al., 2020). At the same time, centromeres in most filamentous fungi are enriched in repetitive DNA, which increases their susceptibility to chromosome breakage and ectopic recombination, making these regions frequent hotspots of chromosomal rearrangements (Dutta et al., 2025; Guin et al., 2020; Smith et al., 2011). However, repeat-poor centromeres have also been reported (Schotanus et al., 2015), suggesting substantial diversity in centromere organization across fungi. Despite their central role in chromosome evolution, centromeric regions have been characterized in only a handful of filamentous fungi (Dutta et al., 2025; King et al., 2015; Schotanus et al., 2015; Seidl et al., 2020; Yadav et al., 2019), leaving the structure, diversity, and evolutionary dynamics of fungal centromeres poorly understood.

In this study, we focus on genome evolution associated with ecological specialization in bark beetle-associated fungi. Bark beetles (Curculionidae: Scolytinae) engage in diverse symbioses with fungi that contribute to their nutrition and host colonization (Biedermann et al., 2019; Hulcr and Stelinski, 2017). Phloem-feeding bark beetles, which acquire most nutrients from plant tissues, are typically associated with facultative fungal symbionts. In contrast, ambrosia beetles bore into nutrient-poor sapwood and rely on obligately cultivated fungal symbionts, termed ambrosia fungi, as their sole food source (Ayres et al., 2000; Batra, 1963; Goodsman et al., 2012; Six, 2012; Six and Elser, 2019). This obligate nutritional symbiosis evolved repeatedly and independently in both beetle and fungal lineages, with no reversals to a non-ambrosial lifestyle reported (Johnson et al., 2018; Jordal and Cognato, 2012; Li et al., 2017; Massoumi Alamouti et al., 2009). These repeated transitions provide a comparative framework for investigating how ecological specialization shapes gene content and genome architecture.

Despite belonging to phylogenetically distant fungal groups, ambrosia fungi display striking phenotypic convergence, including yeast-like growth phases and the production of swollen nutritional cells consumed by beetles (Batra, 1967). However, recent comparative analyses of gene repertoire across phylogenetically independent ambrosia lineages suggests lineage-specific rather than convergent adaptations to the ambrosial lifestyle (Huang et al., 2026). Nevertheless, comparisons across deeply divergent fungal lineages may obscure more subtle evolutionary processes associated with ecological specialization. Importantly, repeated ecological transitions among closely related species provide a unique opportunity to investigate how genome architecture, gene content, and chromosome organization evolve during adaptations. Such questions require chromosome-level genome assemblies that enable analyses of structural genome evolution, including chromosomal rearrangements, centromere evolution, and their relationship to lineage-specific gene evolution.

Here, we investigate genome structure and gene family evolution in *Geosmithia* (Ascomycota: Hypocreales), a widespread and species-rich fungal genus associated with bark beetles (Kolařík and Hulcr, 2023). Throughout its evolutionary history, the genus has given rise to a diversity of ecological lifestyles, making it an outstanding system for studying the genomic basis of adaptation (Kolařík and Hulcr, 2023). Notably, *Geosmithia* has undergone multiple, evolutionarily independent shifts in life strategies, ranging from non-ambrosia to ambrosia symbiosis, from saprotrophic to pathogenic lifestyles, and from generalism to host specialization (Veselská et al., 2019; Zhang et al., 2022). Specifically, three species*, Geosmithia cnesini*, *G. eupagioceri*, and *G. microcorthyli*, represent independent origins of obligate nutritional mutualism with ambrosia beetles (Kolarik and Kirkendall, 2010). Particularly informative is the sister-species pair *G. microcorthyli* and *G. multisociorum*, which differ dramatically in ecological strategy despite their close evolutionary relationship. Whereas *G. microcorthyli* is an obligate ambrosia mutualist exhibiting specialized nutritional adaptations, *G. multisociorum* remains associated with phloem-feeding beetles and lacks these ambrosial characteristics. This contrast provides a rare opportunity to investigate genomic changes accompanying the emergence of obligate fungus farming. In contrast, phytopathogenicity has evolved only once within the genus, represented by *G. morbida*, the causative agent of Thousand Cankers Disease in black walnut (Kolarik et al., 2011; Tisserat et al., 2009). The ecological roles of the remaining *Geosmithia* species are less understood, though they differ markedly in their insect vectors and host plant ranges (Veselská et al., 2019).

Here, we investigate how repeated shifts in ecological strategy have shaped genome evolution in *Geosmithia*. We explore whether transitions in lifestyle are associated with predictable changes in metabolic capacity, gene family composition, chromosome organization, and centromere evolution. To address these questions, we generated chromosome-level genome assemblies and annotations for 11 ecologically diverse *Geosmithia* species using PacBio HiFi sequencing and RNA-seq. In addition, Hi-C data were generated for four representative species, enabling the identification of putative centromeric regions and comparative analyses of centromere organization. Together, these data provide a powerful framework for examining the interplay between chromosome evolution, centromere diversification, and gene repertoire evolution during ecological specialization.

## Results

### Chromosome-level genome assemblies and annotation of ecologically distinct *Geosmithia* species

We first sequenced and assembled 11 *Geosmithia* species including three lineages of ambrosia fungi (*G. cnesini*, *G. eupagioceri*, and *G. microcorthyli*), three pine specialists (*G. abietis* nom. prov.*, G. magnispora*, and *G. pinophila* nom. prov.), and five generalists (*G. multisociorum*, *G. cedri* nom. prov., *G. obscura*, *G. langdonii* and *G. xerotolerans*). Representative genomes from each ecological strategy are shown in **Fig. 1**, with all 11 genomes shown in **Fig. S1**. These species were chosen based on previous phylogenetic analyses suggesting these species represent independent transitions in ecological niche (Zhang et al. 2022). PacBio HiFi sequencing yielded exceptionally high coverage, ranging from 220× to 278×. Near telomere-to-telomere assemblies were obtained for all species, with most chromosomes containing the canonic fungal telomeric repeats (CCCTAA at the 5′ end and TTAGGG at the 3′ end) except for *G. pinophila* nom. prov. and *G. magnispora* with telomere repeats CCCTAAA and TTTAGGG). The main exception involved chromosomes terminating at rDNA loci, which could not be fully assembled through to the telomere resulting in additional short contigs containing rDNA repeats. In *G. abietis* nom. prov.*, G. cnesini*, *G. eupagioceri, G. langdonii, G. magnispora, G. obscura,* and *G. pinophila* nom. prov. one of these short contigs also carried a terminal telomeric repeat, suggesting that the rDNA array is located near a telomeric region in most species, although the precise arrangement of the rDNA repeats and telomere could not be determined for most.

**Figure 1.**
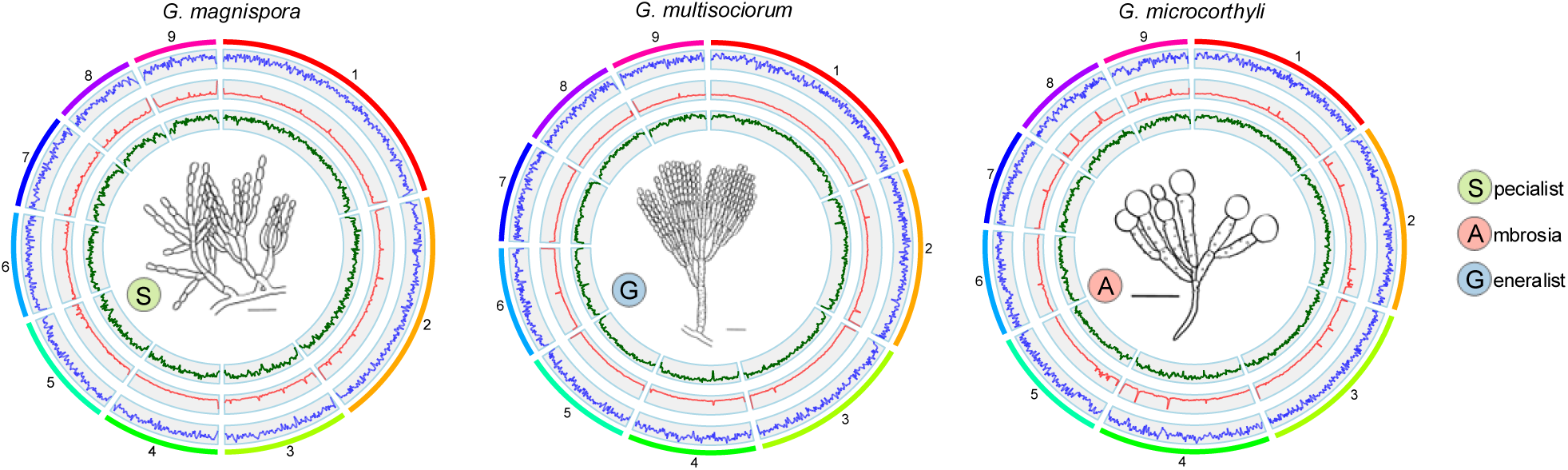
Genome characteristics of representative *Geosmithia* species. Each plot shows genomic features across chromosomes/contigs. Tracks are arranged from the outer to inner circle as follows: numbered chromosomes/contigs, gene density, repeat density, GC content, and illustration of species. Gene density, repeat density, and GC content were calculated in 20 kb windows. Bars indicate 10 µm. Illustrations by Valerie Veselská.

Genomes were assembled into nine contigs representing nine chromosomes in all species except *G. langdonii* and *G. xerotolerans*, in which one chromosome remained unresolved and was represented by two contigs (see section “<u>Extensive genome restructuring across *Geosmithia”* for more details</u>). The assembly correctness was verified by Hi-C sequencing for four species, ambrosial *G. eupagioceri* and *G. microcorthyli*, generalist *G. multisociorum*, and specialist *G. pinophila* nom. prov. Genome sizes ranged from 25.9 Mb in *G. magnispora* to 30.3 Mb in *G. microcorthyli* (**Table 1**).

**Table 1.** Detailed statistics and BUSCO results for genome assemblies and annotation of *Geosmithia* species.

| Species | GS (Mb) | Contigs | Largest contig (Mb) | GC (%) | Busco (%) | Gene number | TE (%) | RIP activity |  |  |
| --- | --- | --- | --- | --- | --- | --- | --- | --- | --- | --- |
|  |  |  |  |  |  |  |  | Affected sequences (%) | LRAR* | Class |
| *G. cedri* nom. prov. (G) | 28.8 | 9 | 5.4 | 53.5 | 94.5 | 8300 | 3.4 | 0.75 | 10 | 2 |
| *G. langdonii* (G) | 28.1 | 10 | 6.3 | 53.6 | 94.9 | 8210 | 3.4 | 0.49 | 8 | 2 |
| *G. multisociorum* (G) | 28.9 | 9 | 5.5 | 53.4 | 94.9 | 8150 | 4.0 | 0.70 | 12 | 2 |
| *G. obscura* (G) | 28.4 | 9 | 5.3 | 52.1 | 94.8 | 8092 | 4.5 | 2.79 | 61 | 3 |
| *G. xerotolerans* (G) | 28.2 | 10 | 5.9 | 53.6 | 95.1 | 8263 | 3.0 | 0.55 | 14 | 2 |
| *G. cnesini* (A) | 26.9 | 9 | 4.6 | 53.8 | 94.4 | 7383 | 6.0 | 0.06 | 0 | 1 |
| *G. eupagioceri* (A) | 29.8 | 9 | 5.7 | 53.4 | 94.5 | 7668 | 7.3 | 0.77 | 0 | 2 |
| *G. microcorthyli* (A) | 30.3 | 9 | 4.7 | 53.4 | 94.6 | 8514 | 4.9 | 0.37 | 0 | 2 |
| *G. abietis* nom. prov. (S) | 28.5 | 9 | 4.6 | 52.6 | 94.2 | 7584 | 11.5 | 0.24 | 0 | 2 |
| *G. magnispora* (S) | 25.9 | 9 | 5.6 | 53.6 | 94.2 | 6993 | 6.1 | 0.03 | 0 | 1 |
| *G. pinophila* nom. prov. (S) | 27.3 | 9 | 6.1 | 52.4 | 94.4 | 7574 | 8.6 | 0.05 | 0 | 1 |

\*Large RIP-Affected Regions: RIP-affected regions spanning at least 4,000 base pairs (bp) (van Wyk et al. 2019). The occurrence and extent of RIP were classified based on (van Wyk et al. 2021) into six classes based on the proportion of RIP affected genome: class 1 –no RIP (0.0 to < 0.2%), Class 2 – low RIP (0.2 to <1.0%), Class 3 –moderately low RIP (1.0 to <5.0%), Class 4 –moderate RIP (5.0 to <10.0%), Class 5 –moderately high RIP (10.0 to <20.0%), and Class 6 high RIP ( ≥ 20.0%). Ecology types: G – generalists, A – ambrosia fungi, S – specialists.

Gene annotation based on transcriptome sequencing and a custom protein database revealed gene number reduction in specialists and ambrosial *G. cnesini* and *G. eupagioceri* compared to generalists (**Table 1**). The number of annotated genes in these species ranged from 6,993 in specialist *G. magnispora* to 7,668 in ambrosial *G. eupagioceri,* whereas gene number in generalists ranged from 8,092 in *G. obscura* to 8,300 in *G. cedri* nom. prov. The genome of ambrosial *G. microcorthyli* bears the largest number of genes, 8,514 (**Table 1**).

The proportion of repetitive elements in genomes was low, ranging from 3.0% in *G. xerotolerans* to 11.5% in *G. abietis* nom. prov. Overall, generalists possessed a lower repeat content (3.0 - 4.5%) than ambrosial species and specialists (4.9 - 11.5%). Single repeats constituted the largest fraction of repetitive elements, accounting for 18 - 73.7% of the total repetitive content across the species. Retroelements represented 0.3 - 53.5% of repetitive elements, whereas DNA transposons were the least abundant category, comprising only 0 - 7.5% of all repetitive elements (**Appendix 1**). GC content was similar across all species with mean value of 53.1% (**Fig. 1, Fig. S1, Table 1**). BUSCO completeness ranged from 94.2% to 95.1% (**Table 1**) and reached similar values as for formerly sequenced *Geosmithia* species (Schuelke et al., 2017).

### No evidence of RIP activity across *Geosmithia*

To evaluate whether the low repeat content of *Geosmithia* genomes results from Repeat-Induced Point mutation (RIP), we analyzed RIP activity across all species. RIP is a fungal genome defense mechanism that limits the proliferation of repetitive DNA, thereby strongly influencing genome evolution (van Wyk et al., 2021). RIP introduces characteristic C:G to T:A mutations into duplicated sequences, thereby inactivating repetitive elements and contributing to AT-rich genomic regions (Galagan and Selker, 2004). Because RIP operates specifically during meiosis, its presence is often considered indirect evidence of sexual reproduction in fungi otherwise regarded as asexual. Overall, our analyses revealed little or no evidence of RIP activity across *Geosmithia* genomes (**Table 1**). The highest proportion of RIP-affected regions was detected in *G. obscura* (2.79%); however, this value still corresponds to RIP class 3, indicating only moderately low RIP activity. These findings support the predominantly asexual nature of *Geosmithia*. Interestingly, pine-specialist species, which possess relatively higher proportions of repetitive DNA, exhibited the lowest RIP activity, suggesting that reduced RIP efficiency may contribute to additional repeat accumulation in these lineages.

### Repeated evolution of specialization and ambrosial state

Comparing species that have independently evolved similar adaptive traits is a powerful approach for identifying the genetic and genomic underpinnings of those traits. Phylogenetic relationships within *Geosmithia* have previously been inferred using only four loci: ITS, TEF1-α, TUB2, and RPB2 (Aylward et al., 2024; Huang et al., 2017; Zhang et al., 2022), providing limited resolution of genome-wide evolutionary relationships. To obtain a robust phylogenomic framework for our comparative analyses, we reconstructed the species phylogeny from 3,816 single-copy orthologous BUSCO genes (**Fig. 2**). The resulting topology was found to be congruent with the previously published phylogenies (Zhang et al., 2022), confirming the independent origins of ambrosial and pine specialist lineages within the genus (**Fig. 2**). Interestingly, low gene and site concordance factors at several internal nodes indicate substantial gene-tree discordance in *Geosmithia*, consistent with incomplete lineage sorting for numerous BUSCO loci.

**Figure 2.**
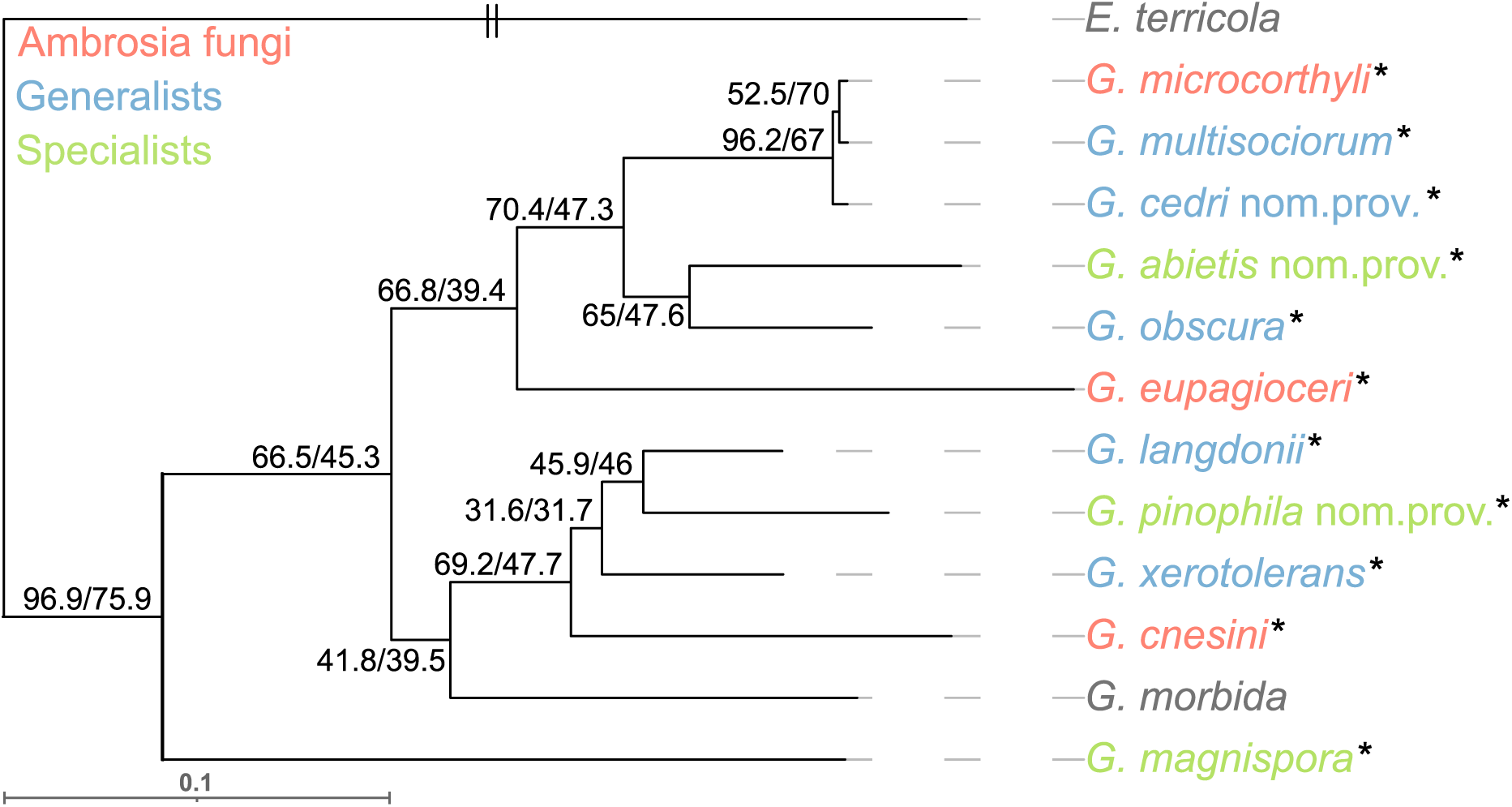
Phylogenomic relationships of *Geosmithia* showing independent origins of ecological strategies. Phylogeny was inferred from a concatenated alignment of 3,816 single-copy orthologous BUSCO genes using maximum likelihood in IQ-TREE. Numbers at nodes indicate gene concordance factor (gCF) and site concordance factor (sCF), representing the percentage of decisive gene trees and alignment sites, respectively, that support a given bipartition. The branch marked with // was shortened to one-half of its original length for visualization. Species sequenced in this study are marked with an asterisk and color-coded by ecology: ambrosia symbionts (red), generalists (blue), specialists (green). *Geosmithia morbida* is a well-known plant pathogen. The genomes of *G. morbida* and *Emericellopsis terricola* (outgroup) were downloaded from NCBI.

### Specialization leads to metabolic streamlining

Our genome annotations revealed gene number reductions in pine specialists and two ambrosia fungi – *G. cnesini* and *G. eupagioceri –* compared to generalists. To assess how gene reduction affects metabolic ability, we first annotated gene functions using eggNOG-mapper, which provides KEGG (Kyoto Encyclopedia of Genes and Genomes) orthology identifiers for genes, followed by specialized annotation pipelines targeting key functional categories. EggNOG-mapper assigned functional annotations to 80.0 - 87.7% of predicted genes across the analyzed genomes, of which 48.0 - 53.2% were assigned at least one function based on the KEGG Orthology (KO) database (**Table 2**). In total, we identified 3,346 unique KOs across the dataset, of which 3,133 were shared by all species, indicating a largely conserved core metabolic repertoire in *Geosmithia*. KEGG pathway reconstruction suggests that all species possess the complete gene sets required for the biosynthesis of all amino acids (**Fig. S2**), as well as for the synthesis of vitamin B6, riboflavin, and thiamine (**Fig. S3**). The most pronounced functional difference was a consistent reduction in gene numbers across all annotated categories in pine specialists compared to generalists (**Table 2**, **Fig. 3**). However, this difference was slightly under statistical significance (p-values: Carbohydrate-Active enZYmes (CAZymes) – 0.054, signal proteins – 0.051, pathogen-host interaction (PHI) analysis – 0.058). Among ambrosial species, *G. cnesini* and *G. eupagioceri* exhibited gene counts comparable to pine specialists, whereas *G. microcorthyli* more closely resembled its sister lineage (*G. multisociorum* and *G. cedri* nom. prov.), likely reflecting the relatively short divergence time between these two lineages.

**Figure 3.**
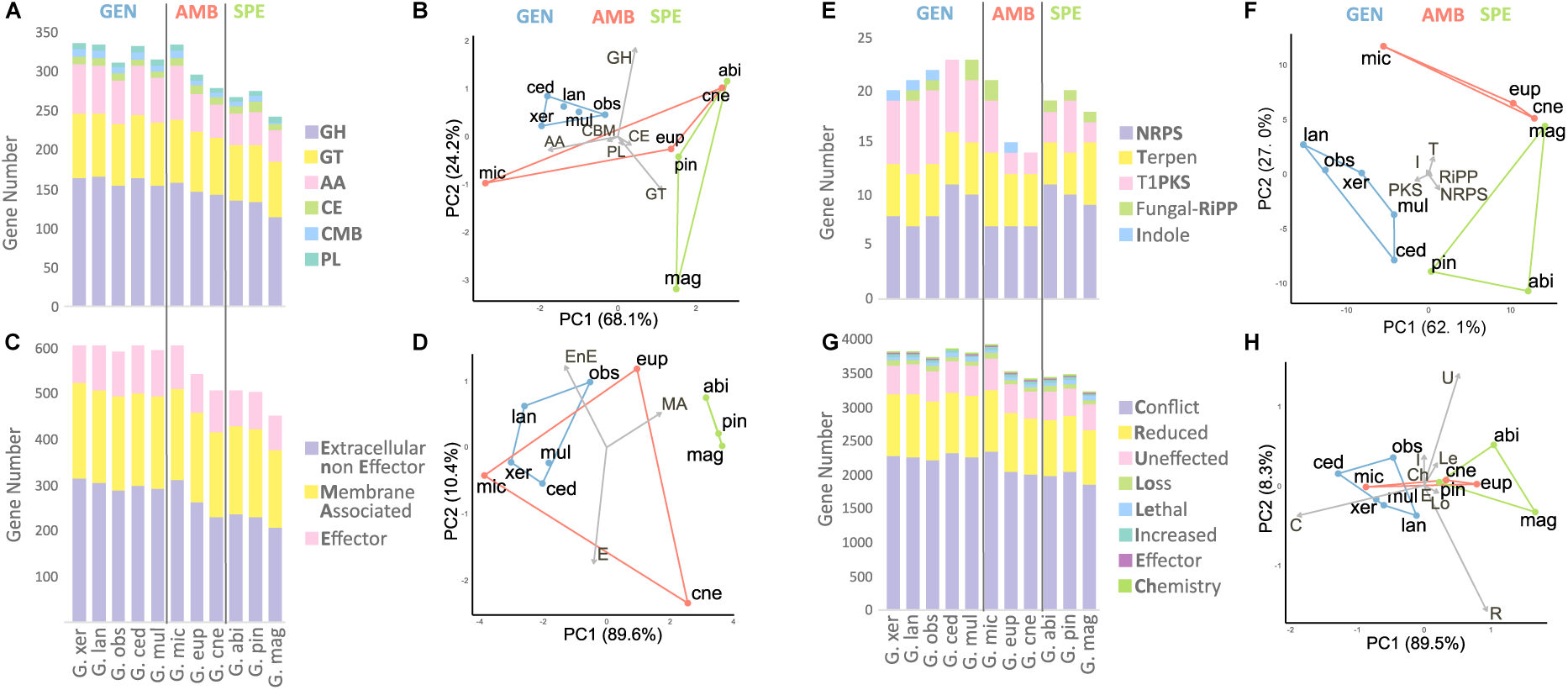
Specialization is accompanied by gene losses in *Geosmithia*. Stacked bar plots show absolute numbers of genes assigned to individual functional categories in each genome: (A) CAZymes: GH – Glycoside Hydrolases, GT – Glycosyl Transferases, AA – Auxiliary Activities, CE – Carbohydrate Esterases, CMB –Carbohydrate-Binding Modules, PL – Polysaccharide Lyases and (B) PCA. (C) Signal proteins and (D) PCA. (E) Secondary metabolites, and (F) PCA. (G) Genes involved in PHI: pathogen – host interactions and (H) PCA. Principal component analyses (PCA) were performed using the proportional representation of each class within individual functional groups. Species are colored according to ecology: generalists (blue), ambrosia symbionts (red), and specialists (green). Percentages on axes indicate the variance explained by each principal component.

**Table 2.**
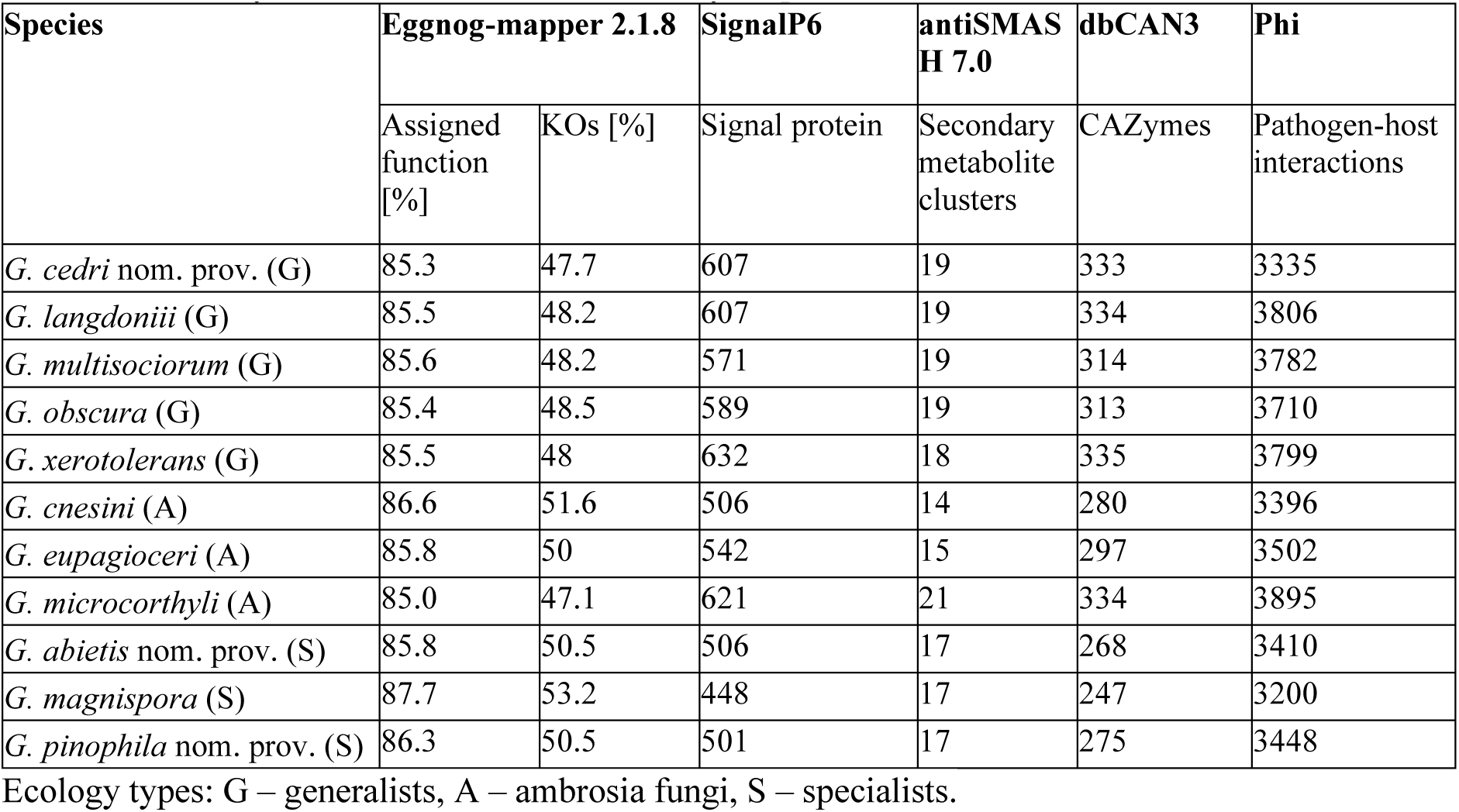
Summary of results from functional analyses.

Compared to generalists, pine specialists exhibited a pronounced contraction of their CAZyme repertoire (**Fig. 3A, B**; **Table S1**). Specifically, they lacked several subfamilies belonging to auxiliary activities (AA): AA13, Carbohydrate Esterases (CE): CE8 and Glycoside Hydrolases (GH): GH114, GH31_13, GH31_2, and GH43_33, and showed reduced copy numbers in AA7, GH13_1, GH18, GH188, GH3, and Glycosyl Transferases GT2 (**Appendix 1**). These families are primarily associated with the degradation of starch, hemicellulose, and chitin, indicating a streamlining of plant and fungal cell wall and storage polysaccharide degradation capacity in specialist lineages. Despite this contraction, complete KEGG pathways for the degradation of these substrates remain encoded in specialist genomes, suggesting that gene loss predominantly affects redundancy and enzymatic diversity rather than eliminating core metabolic functionality.

Patterns in signal proteins further support this specialization driven genome streamlining (**Fig. 3C, D; Table S2**). While the number of membrane-associated signal proteins was comparable across species, pine specialists showed a marked reduction in predicted extracellular non-effector proteins and candidate effectors relative to generalists. Because secreted proteins mediate host interaction and environmental substrate exploitation, their contraction is consistent with reduced ecological breadth and host-range restriction. Secondary metabolite repertoires exhibited a similar trend (**Fig. 3E, F; Table S3**). Pine specialists possessed fewer type I polyketide synthase (T1PKS) clusters, but an expansion of NRPS-like clusters compared to generalists. This shift may reflect a reconfiguration of chemical ecology, potentially favoring peptide-based metabolites over polyketide diversity in specialized host environments. Genes with predicted roles in pathogen-host interactions were reduced in pine specialists and ambrosial *G. cnesini* and *G. eupagioceri,* especially in groups classified as conflict (with more than one definition of phenotype), reduced, and unaffected (**Fig. 3G, H**; **Table S4**).

### Gene family evolution

To explore whether ecological shifts in *Geosmithia* were associated with specific gene family loss or expansion, we compared all annotated genes across the 11 *Geosmithia* genomes by first assigning them to orthogroups (OGs). We identified 8,616 orthogroups across the analyzed *Geosmithia* genomes, of which 5,912 were shared by all species and 376 were identified to be under positive selection in combined BUSTED/aBSREL analyses in Hyphy (**Appendix 1**). The vast majority of OGs were present as single-copy genes, with proportions ranging from 88.9% to 94.5% across species (**Table S5**). Only a small fraction (4 - 10.4%) were multicopy, and species-specific OGs were rare, typically comprising only a few genes per genome (0 - 65). Interestingly, expanded and species-specific gene families were predominantly associated with repeat-rich or subtelomeric regions, while genes under positive selection were more broadly distributed across chromosomes (**Fig. 4A**, **Fig. S4**). The highest number of species-specific OGs were present in the genome of the ambrosial species *G. microcorthyli*. Several species did not possess any species-specific OGs including *G. magnispora* (specialist), *G. cedri* nom. prov. (generalist), and *G. multisociorum* (generalist).

**Figure 4.**
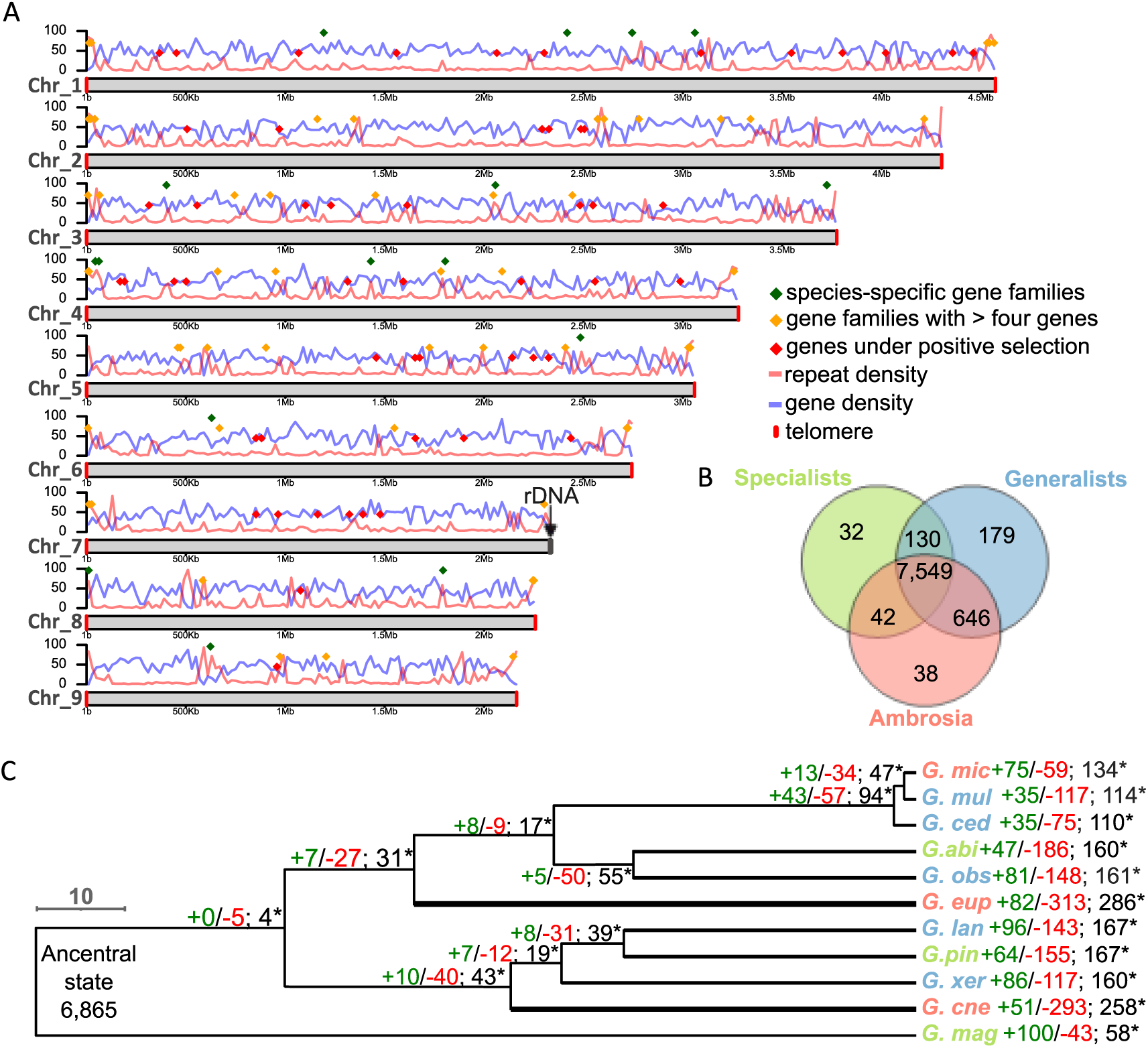
Gene family evolution in *Geosmithia*. (A) Expanded and species-specific gene families localize close to repetitive and subtelomeric regions. Representative genomic landscape is shown for *G. abietis* (specialist). Blue line represents gene density, and red line repeat density. Colored diamonds mark the genomic positions of genes from different evolutionary categories: genes under positive selection (red), gene families with > 4 copies (yellow), and species-specific gene families (green). Chromosome ends marked in red denote telomeric regions, whereas grey regions indicate rDNA loci. (B) Orthogroups shared across ecological groups in *Geosmithia*. An orthogroup was considered present in a given ecological group when it occurred in at least one species belonging to that group. (C) Gene family evolution. Red values indicate gene family contractions (losses) and green values indicate expansions (gains) along each branch. Branch width is proportional to the number of significantly evolving orthogroups detected on that lineage.

Comparison of OG composition across ecological groups showed that most were shared among multiple lifestyles, indicating a substantial conserved core gene repertoire within the genus (**Fig. 4B**). At the same time, each ecological group retained a subset of unique or preferentially represented OGs, consistent with lineage-specific gene content differentiation accompanying ecological specialization. We found no clear association between ecological strategy and the identity of genes evolving under positive selection (**Appendix 1**). Across all species, 376 orthogroups showed signatures of positive selection. Of these, 196 (52%) received functional annotations using our pipeline. Functionally annotated genes included 10 CAZymes, 7 candidate effectors, 3 secondary metabolite genes, and 178 genes associated with pathogen–host interactions, although some genes belonged to multiple functional categories.

To explore how gene family evolution contributes to ecological transitions, we used CAFE to infer gene family expansions and contractions across the reconstructed species phylogeny based on 6,554 orthogroups (OGs) shared by at least two species (**Fig. 4C**). The reconstructed last common ancestor of *Geosmithia* was estimated to contain 6,865 gene copies across the modeled orthogroups. Changes in gene family size occurred predominantly along terminal branches, with contractions generally exceeding expansions, except in a specialist *G. magnispora* and ambrosial fungus *G. microcorthyli*, where expansions prevailed. Branches with the greatest widths, corresponding to the highest numbers of significantly evolving OGs, highlight clades that experienced particularly dynamic gene family evolution. In total, 768 gene families showed significantly accelerated evolutionary rates, of which 636 were significant in more than one node. Among these rapidly evolving families, 44 were assigned to CAZymes, representing 13.5% of all annotated CAZyme OGs; 402 to PHI-base-associated families (9.9% of all PHI-annotated OGs); 22 corresponded to effector families (12.9% of all effector OGs); 84 to secreted non-effector proteins (37.2% of all signal peptide-containing non-effector OGs); and 44 to secondary metabolite-associated families (21.6% of all annotated secondary metabolite OGs). Within this category, gene family encoding NRPS-like SM was significantly expanded in pine specialists, whereas the family encoding fungal-RiPP-like SM was significantly decreased and one of NRPS families absent in ambrosia fungi. Overall, 40.5% of significantly evolving gene families were not assigned to any of the analyzed functional categories, similar to the proportion observed across all orthogroups (47.3%) (**Appendix 1**).

### Atypical centromere structure and evolution in *Geosmithia*

Beyond gene content, chromosome-level assemblies enabled us to investigate the evolution of genome architecture. We therefore examined centromere organization, chromosome structure, and large-scale genome rearrangement across *Geosmithia*. First, Hi-C contact maps were generated for four *Geosmithia* species representing all three lifestyles to verify our genome assemblies. After detailed inspection, we identified Hi-C contacts consistent with a Rabl chromosome configuration, (**Fig. 5A**, **Fig. S5**) characterized by clustering of centromeres on one side of the nucleus and telomeres on the opposite side (Torres et al., 2023). Because these regions lack extensive repetitive DNA, the observed contacts cannot be explained by ambiguous read mapping and therefore provide strong evidence for true centromere clustering. Centromeric positions were consistently identified from inter-centromeric contact dots visible at shared centromeric coordinates (**Fig. 5A**; **Fig. S5**). The inferred centromeres indicate that *Geosmithia* possesses an unusual centromere architecture compared to other fungi. The centromeric regions are gene-rich, contain very low repeat content, and harbor numerous actively transcribed genes (**Fig. 5B**). Moreover, these loci form conserved syntenic blocks easily identified across species, suggesting long-term stability during genome evolution and maintenance of chromosome number (n = 9) (**Fig. 5C**).

**Figure 5.**
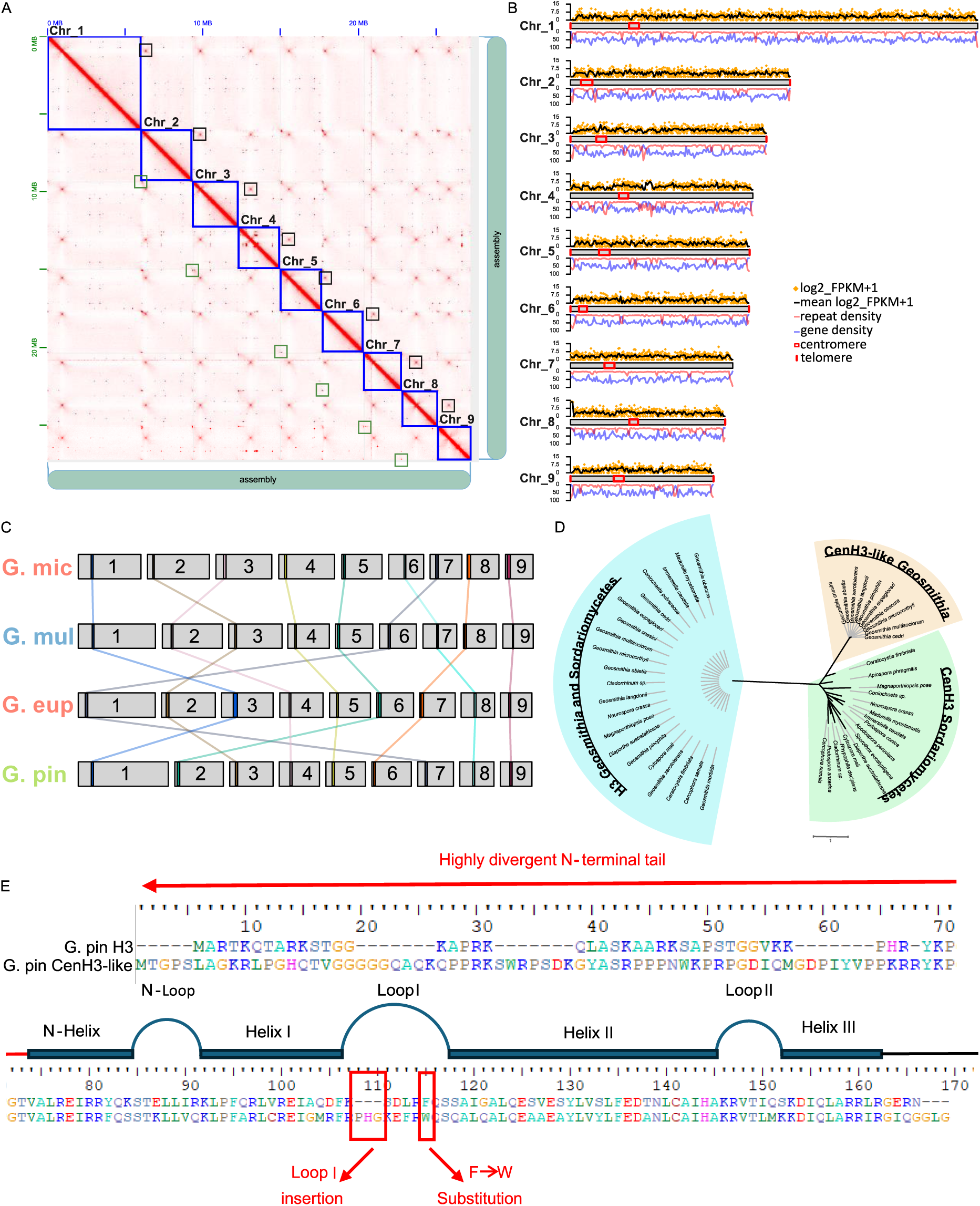
Centromere organization and evolution in *Geosmithia*. (A) Hi-C contact map illustrating chromosome-scale organization and centromere and telomere clustering (Rabl configuration) in *G. pinophila* nom. prov. genome. Blue squares highlight chromosomes (assembled as contigs), while focal high-interaction regions correspond to putative centromeres. Black squares indicate inter centromeric interactions while green squares indicate inter telomeric contacts. (B) Genomic profiles along chromosomes showing high gene expression (log₂ FPKM + 1 in orange, mean expression in 20 kb windows in black), high gene density (blue), and low repeat density (red) in centromeric regions (red rectangles). (C) Conserved syntenic centromeric gene blocks across four Hi-C-sequenced *Geosmithia* species, illustrating that chromosome numbering based on size does not correspond to chromosome homology. (D) Phylogenetic analysis of histone H3 and CenH3 proteins across *Geosmithia* and representative Sordariomycetes. *Geosmithia* species lack canonical CenH3 and instead possess a divergent CenH3-like variant forming a distinct clade separate from conserved fungal CenH3 homologs. (E) Protein alignment highlighting the major structural differences between canonical histone H3 and the putative CenH3-like protein identified in *Geosmithia*. The CenH3-like protein is characterized by (i) a highly divergent N-terminal tail, (ii) an insertion within Loop I of the histone-fold domain, and (iii) substitution of the highly conserved phenylalanine residue by tryptophan (F→W). The positions of the N-helix, α-helices (Helices I–III), and connecting loops are indicated according to the canonical histone H3 structure.

Positions of centromeres in filamentous fungi are specified epigenetically by the centromere-specific histone CenH3 (also known as CENP-A or Cse4), a divergent variant of canonical histone H3 (Dutta et al., 2025; Guin et al., 2020; Smith et al., 2011). Searches of annotated proteomes recovered two proteins annotated as histone H3 in *Geosmithia* species, in addition to the remaining core histones H2A/H2B/H4. No protein was annotated as a CenH3 homolog. To determine the identity of the two H3 proteins, we performed phylogenetic analyses together with canonical H3 and characterized CenH3 proteins from representative Sordariomycetes. The resulted phylogeny clearly separated canonical H3 and CenH3 proteins and confirmed that one of the two *Geosmithia* H3 proteins clustered with canonical H3 (**Fig. 5D**). All previously characterized CenH3 proteins from Sordariomycetes were successfully recovered by the HMM pipeline, demonstrating its sensitivity. In contrast, the second *Geosmithia* H3 protein formed a distinct CenH3-like lineage together with a homolog from *G. morbida* and was clearly separated from both canonical H3 and previously characterized CenH3 proteins (**Fig. 5D**). The only exception was *G. magnispora*, in which no CenH3-like protein was detected. This CenH3-like protein is a strong candidate for a highly divergent centromere-specific histone variant. It carries sequence features associated with centromeric histones (**Fig. 5E**), including a highly divergent N-terminal tail and modified loop1 by an insertion and substitution of highly conserved phenylalanine by a tryptophan (Keith et al., 1999). Re-analysis of genome derived ORFs, including unplaced contigs, again failed to detect any conventional CenH3 sequence, supporting that the absence of typical CenH3 in *Geosmithia* is biological rather than technical.

### Extensive genome restructuring across *Geosmithia*

The varying chromosomal position of the nine conserved centromeric regions suggested substantial genomic rearrangement across *Geosmithia*. To investigate chromosomal evolution within the genus, we examined large-scale rearrangements using synteny analyses implemented in GENESPACE (**Fig. 6**). These analyses revealed that intra and interchromosomal rearrangements are widespread across *Geosmithia*. Despite extensive chromosomal reshuffling, all analyzed *Geosmithia* species retained a conserved karyotype of nine chromosomes which appear to be specified by the nine unique centromere blocks. The only exceptions were *G. xerotolerans* and *G. langdonii*, which were assembled into 10 contigs, likely the result of challenges of assembling across the rDNA locus. Synteny analyses, together with the conserved position of the terminal rDNA locus across the genus, indicate that the shortest contig in *G. langdonii* represents a fragment of chromosome 8, whereas the shortest contig in *G. xerotolerans* most likely corresponds to a fragment of chromosome 1.

**Figure 6.**
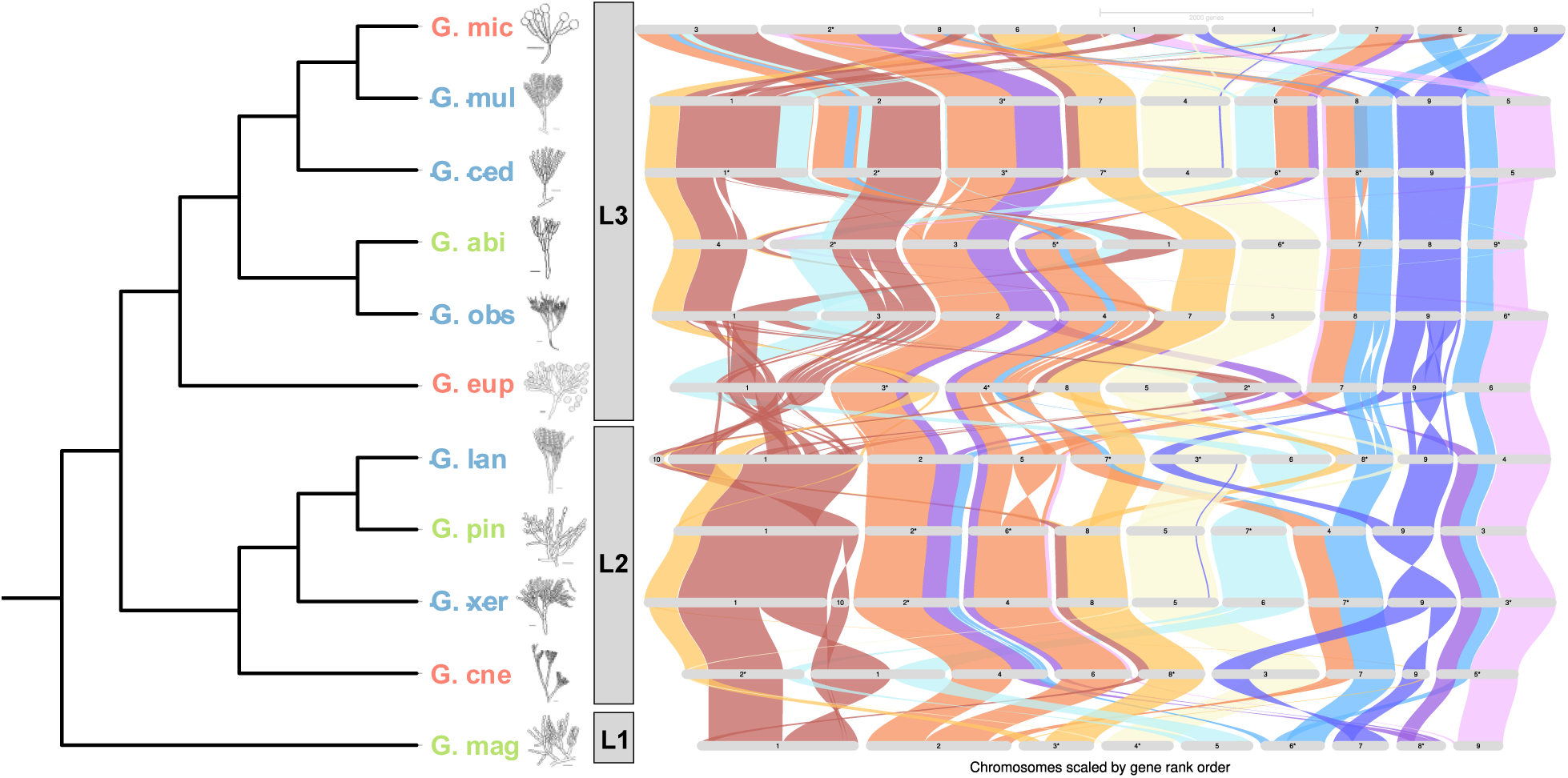
Extensive chromosomal rearrangement in *Geosmithia*. Genome-wide synteny visualized using GENESPACE. Colored ribbons connect orthologous gene blocks across genomes, with *G. magnispora* used as the reference genome. Species are divided into three lineages: L1 (*G. magnispora),* L2 (*G. cnesini, G. xerotolerans, G. pinophila* nom. prov., *G. langdonii*), L3 (*G. eupagioceri, G. obscura, G. abietis* nom. prov., *G. cedri* nom. prov., *G. multisociorum, G. microcorthyli*).

Synteny analyses revealed contrasting patterns of chromosomal evolution across *Geosmithia.* The largest chromosomal changes were observed between the three major *Geosmithia* clades as illustrated by the extensive rearrangements in the synteny comparisons between *G. langdonii* and *G. eupagioceri* and between *G. magnispora* and *G. cnesini* (**Fig. 6**). Comparisons between species revealed that the rate of chromosomal changes do not appear uniform across the genus. *G. pinophila* nom. prov., and *G. xerotolerans* retained highly similar chromosome organization while contrasts between *G. pinophila* and its sister species, *G. langdonii* show extensive rearrangement. Likewise, the closely related generalists *G. cedri* nom. prov. and *G. multisociorum* exhibited near perfect synteny while the ambrosial species *G. microcorthyli* displayed extensive chromosome restructuring despite sharing approximately 99% genome-wide sequence identity with its sister species *G. multisociorum* (**Fig. 6**, **Fig. 7B**). These observations indicate that while extensive chromosomal differences are generally associated with deeper phylogenetic contrasts, large scale chromosomal changes can also accumulate rapidly over very short evolutionary timescales.

**Figure 7.**
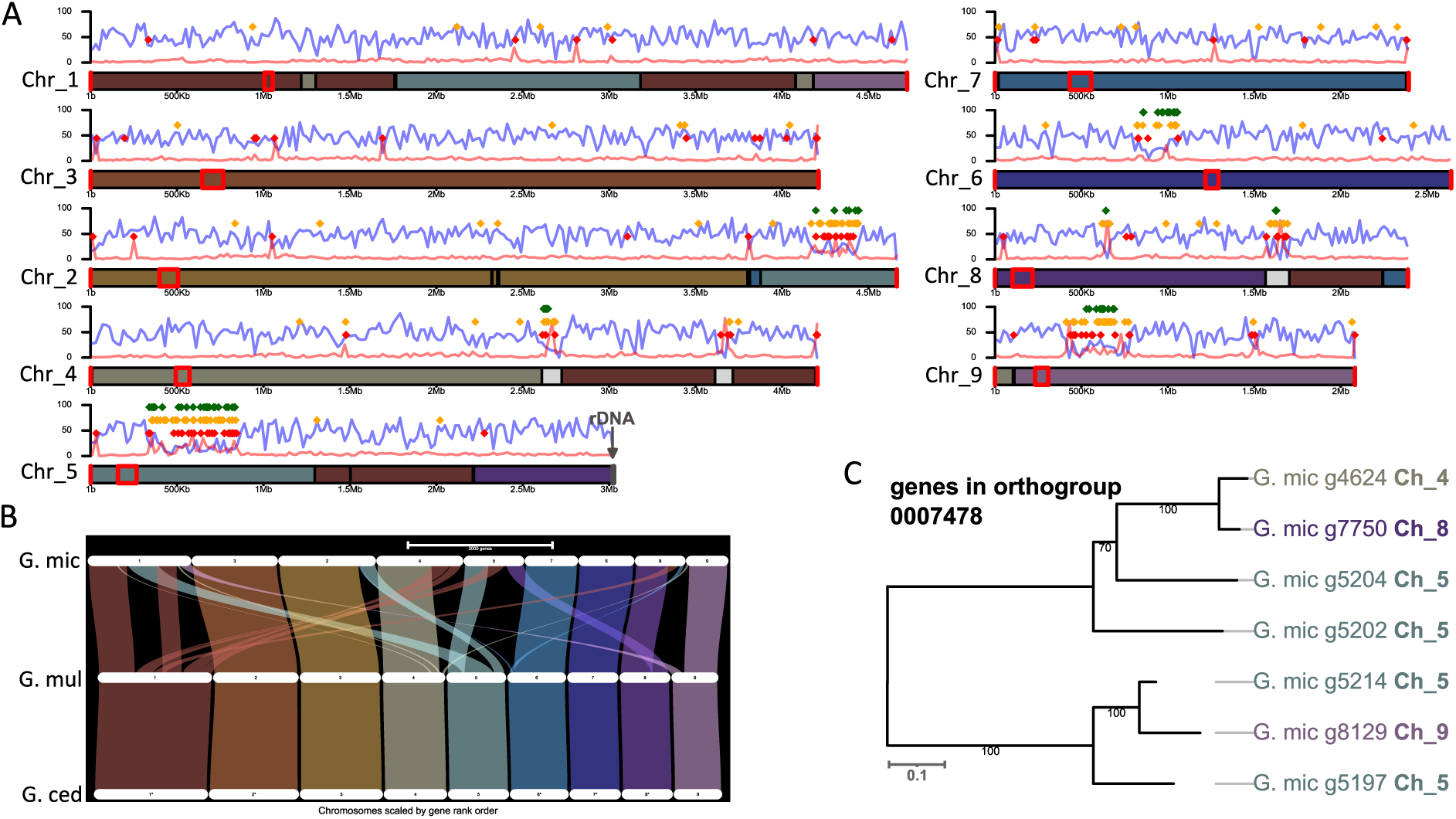
Unique genome architecture of *G. microcorthyli*. (A) KaryoploteR visualization of gene (blue) and repeat (red) densities along the chromosomes of *G. microcorthyli*. Colored diamonds indicate LTR/Gypsy retrotransposons (red), gene families containing at least five members (yellow), and species-specific gene families (green). Chromosome ends highlighted in red denote telomeric regions, whereas grey regions indicate rDNA locus. Red rectangles on the chromosome ideograms indicate the putative positions of centromeres inferred from the Hi-C contact map. Colored segments of the ideograms represent syntenic blocks identified by GENESPACE between *G. microcorthyli* and its sister species, while grey segments indicate regions specific to *G. microcorthyli*. (B) Syntenic relationships between *G. microcorthyli* and its sister species identified by GENESPACE. (C) Maximum-likelihood phylogenetic tree of one representative orthogroup (OG0007478) illustrating the distribution of paralogous genes across the chromosomes of *G. microcorthyli*. Tip labels are color-coded according to the corresponding syntenic blocks identified by GENESPACE.

### Ambrosial *G. microcothyli* displays unique genome architecture among *Geosmithia*

To further characterize the exceptional genome organization of the ambrosial species *G. microcorthyli* (**Fig. 7B**), we examined the distribution of chromosomal breakpoints and other genomic features along its chromosomes (**Fig. 7A**). Chromosomal breakpoints frequently occurred within gene-dense regions rather than within repeat-rich compartments, e.g. on chromosome 1, 5 or 9. In contrast, repetitive DNA was concentrated in several discrete chromosomal islands dominated by Gypsy retrotransposons. Such prominent repeat-rich islands were not observed in the remaining *Geosmithia* species. These repeat islands also harbored numerous species-specific genes and expanded gene families. Consistent with this pattern, *G. microcorthyli* possessed the highest number of species-specific orthogroups among all analyzed species (**Table S5**). Nine of twenty species-specific othogroups were functionally annotated using our pipeline, and all were assigned to the pathogen-host interaction category (Supplementary Material 1). Notably, members of expanded orthogroups were distributed across multiple chromosomes rather than arranged as tandem arrays within individual repeat-rich islands (**Fig. 7C**).

## Discussion

Comparative genomics has revealed extensive variation in genome architecture both within fungal species and genera (Fouché et al., 2026; Li et al., 2023; Quenu et al., 2022; Shi-Kunne et al., 2018) and structural genome evolution it is increasingly recognized as a major driver of genome diversification, adaptation, and speciation in fungi (Rajeh et al., 2018). However, despite rapid progress in chromosome-scale fungal genomics, current understanding of structural genome evolution remains strongly biased toward plant pathogens and yeasts (Ament-Velasquez et al., 2024; Ma et al., 2010; Quenu et al., 2022; Schotanus et al., 2015; Shi-Kunne et al., 2018; Treindl et al., 2021). Here we demonstrate extensive chromosomal rearrangements within a symbiotic genus *Geosmithia* providing insight into genome evolution associated with repeated and evolutionarily independent ecological transitions, including shifts toward ambrosia symbiosis and host specialization. Synteny analyses of our genome assemblies revealed pervasive intra- and inter-chromosomal rearrangements across eleven *Geosmithia* species. The synteny analyses reveal two contrasting but non-exclusive evolutionary scenarios. In the first, chromosomal architecture appears to remain stable over extended evolutionary periods, as illustrated by the near-perfect synteny between *G. xerotolerans* and *G. pinophila* nom. prov. In the second scenario, extensive chromosomal reshuffling can occur over very short evolutionary timescale, as evidenced by the low synteny between the sister species *G. microcorthyli* and *G. multisociorum*, despite their extremely high whole-genome sequence similarity (∼99%). Together, these patterns suggest that chromosomal evolution in *Geosmithia* may be characterized by bursts of rapid genome restructuring followed by long periods of structural stasis, rather than a constant rate of rearrangement. In fungi, stressful environmental conditions have been shown to promote genome structural variability, which can subsequently generate phenotypic variation with adaptive potential (Vande Zande et al., 2023). Although the present data does not allow us to conclude whether chromosomal rearrangements in *Geosmithia* are adaptive, we hypothesize that may have been triggered by ecological transitions associated with novel environmental conditions, such as shifts in host tree chemistry, nutritional environment, or interactions with new beetle vectors.

Interestingly, chromosomal breakpoints in *Geosmithia* were not associated with repeat-rich regions as it was described in plant pathogenic fungi in which TE proliferation is considered a major driver of genome structural alternations (Alkemade et al., 2025; Fouché et al., 2026; Ma et al., 2010; Priest et al., 2020; Shi-Kunne et al., 2018). Instead, rearrangements frequently occurred within gene-dense genomic regions, suggesting that alternative mechanisms may contribute to chromosome instability in compact fungal genomes. One possible explanation involves non-B DNA-forming sequence motifs, which alternate 3D DNA conformation and promote double-strand breaks and genome structural instability (Zhao et al., 2010). Although analyses of non-B DNA motifs were beyond the scope of the present study, our results indicate that TE-independent mechanisms may represent an important and currently overlooked driver of structural genome evolution in fungi.

The low abundance of repetitive DNA in *Geosmithia* is also reflected in its atypical centromere organization. In most filamentous fungi, centromeres are repeat-rich, heterochromatic regions specified epigenetically by the centromeric histone CenH3 (also termed CENP-A or Cse4p) and are considered among the fastest evolving genomic regions (Dutta et al., 2025; Seidl et al., 2020; Smith et al., 2011). Their evolutionary plasticity, including centromere repositioning, inactivation, and neocentromere formation, has been implicated in karyotype evolution by stabilizing newly rearranged chromosomes and facilitating chromosome fusions and chromosome-number changes (Guin et al., 2020; Narayanan et al., 2024; Sankaranarayanan et al., 2020). In contrast, centromeres in *Geosmithia* are not highly repetitive and instead harbor transcriptionally active genes and form conserved syntenic blocks easily identified across species. Such conservation contrasts with the prevailing view of rapidly evolving fungal centromeres and suggests that centromere positions in *Geosmithia* remained evolutionarily stable despite extensive chromosome reshuffling occurring along chromosomal arms. We hypothesize that this exceptional centromeric stability may have contributed to the maintenance of a conserved nine-chromosome karyotype despite extensive structural genome evolution.

Interestingly, despite low repeat content and presence of transcribed genes, putative centromeric regions displayed characteristic Hi-C interaction patterns similar to those described for repeat-rich fungal centromeres. These regions formed focal interaction hotspots within a chromosome and extensive inter-centromeric contacts between chromosomes consistent with a classical Rabl chromosome configuration (Torres et al., 2023), in which centromeres cluster at one pole of the nucleus while telomeres localize to the opposite pole. Together, these observations emphasize how little is still known about centromere evolution in filamentous fungi, for which only a small number of species have been characterized in detail (Dutta et al., 2025; King et al., 2015; Seidl et al., 2020; Smith et al., 2011; Yadav et al., 2019).

Our analyses further suggest that centromere identity in *Geosmithia* may be specified by a highly divergent CenH3-like protein that forms a distinct lineage separate from CenH3 proteins of other Sordariomycetes. Importantly, none of the annotated CenH3 proteins included in our dataset originated from species belonging to the same family (Bionectriaceae) as *Geosmithia*, limiting direct phylogenetic comparisons with closely related taxa. Nevertheless, the identified CenH3-like protein in *Geosmithia* contains several characteristic features associated with functional CenH3 proteins, including a highly divergent N-terminal tail and sequence changes within the conserved loop 1 region, including an insertion and substitution of the highly conserved phenylalanine by tryptophan (Keith et al., 1999). These observations suggest that CenH3 proteins may display considerably greater sequence diversity than previously appreciated. However, functional validation will be necessary to confirm the role of this CenH3-like protein in centromere determination.

Despite the overall low repeat content, its proportion differed substantially among ecological groups. Generalists exhibited the lowest repeat content and at the same time stronger signatures of RIP, including the presence of LRAR, which was absent in specialists and ambrosia fungi. As RIP operates specifically during the sexual cycle (Galagan and Selker, 2004), these patterns may reflect differences in the evolution of reproductive strategies among ecological groups. The stronger RIP signatures detected in generalists may indicate prolonged retention of sexual reproduction, whereas reduced RIP activity in specialists and ambrosia fungi could reflect their earlier transitions toward predominantly clonal lifestyles. Besides lower RIP activity, higher proportion of repeat content in specialists and ambrosia fungi may be facilitated by reductions in effective population size associated with ecological specialization. Pine specialists are restricted to a narrow range of vectors and host trees (Kolařík and Hulcr 2023) and ambrosia fungi are tightly associated with individual beetle lineages and vertically transmitted by their hosts (Hulcr and Stelinski, 2017; Six and Wingfield, 2011), which in both systems may result in fragmented populations. Smaller effective population size is related to relaxed purifying selection and intensification of the effects of random genetic drift in genome evolution (Lynch and Conery, 2003). These processes could result in gradual accumulation of repetitive sequences in genomes as we see in *Geosmithia* specialists and ambrosia species. Although the accumulation of repetitive elements could be a result of random processes, it does not mean that they do not have an adaptive outcome (Lynch and Conery, 2003). Transposable elements were indicated as a tool for generating genetic variability in asexual fungi (Alkemade et al., 2025; Dong et al., 2015; Torres et al., 2020). Consistent with this hypothesis, expanded and species-specific gene families in *Geosmithia* were preferentially associated with repeat-rich regions, suggesting that localized repeat accumulation may create genomic environments permissive for gene innovation and diversification. This pattern was particularly pronounced in the ambrosia species *G. microcorthyli*, whose chromosomes contain discrete repetitive islands enriched in Gypsy retrotransposons and harboring nested species-specific genes. Notably, genes belonging to the same orthogroup were dispersed across multiple chromosomes rather than occurring as tandem duplications. We hypothesize that the activity of Gypsy retrotransposons facilitated this distribution by mobilizing adjacent genomic sequences during transposition, thereby promoting the spread of these genes among chromosomes.

Although we detected species specific gene family expansions, gene loss appears to have been the dominant process accompanying ecological specialization in *Geosmithia*. Substantial gene reduction was observed in pine specialists and in the ambrosia species *G. cnesini* and *G. eupagioceri*, affecting all analyzed functional categories. Nevertheless, core metabolic pathways involved in plant cell wall degradation and amino acid biosynthesis remained conserved across all species. These observations suggest that ecological specialization in *Geosmithia* may involve metabolic streamlining through preferential loss of redundant genes rather than complete elimination of metabolic pathways. This pattern is consistent with functional minimization accompanying ecological specialization (Baroncelli et al., 2016; Shi-Kunne et al., 2018; Veselská et al., 2020), where adaptation to a narrower and chemically more homogeneous host environment reduces the selective advantage of maintaining a broad degradative arsenal. Whether these gene losses are preferentially associated with chromosomal breakpoints remains unresolved and represents an important direction for future research.

Notably, predicted metabolic capabilities did not always correspond to experimentally observed phenotypes, emphasizing that the presence of genes alone may not accurately predict functional activity (Kjærbølling et al., 2020). Previous phenotypic microarray analyses and enzymatic assays in *Geosmithia* revealed reduced metabolic versatility in pine specialists and, to a lesser extent, in ambrosia species (Veselská et al., 2019). This pattern is further supported by the inability of pine specialists to grow in the absence of essential nutrients in cultivation media, suggesting increased nutritional dependence associated with ecological specialization (Kolařík and Hulcr, 2023).

To conclude, our results challenge the prevailing view that extensive structural genome evolution in fungi is primarily driven by transposable element proliferation and rapidly evolving centromeres. Instead, *Geosmithia* demonstrates that extensive chromosome restructuring can occur in compact, gene-dense genomes with low repeat content and evolutionarily stable centromeres. The striking contrast between the recently diverged sister species *G. multisociorum* and *G. microcorthyli*, which share nearly identical genome sequences yet differ markedly in chromosome architecture, gene content, and repetitive DNA organization, provides a unique opportunity to investigate the mechanisms underlying structural genome evolution during ecological diversification. Together, these findings establish *Geosmithia* as a powerful model system for linking chromosome evolution, gene turnover, and ecological adaptation, and open new avenues for exploring the roles of chromosomal breakpoints, transposable elements, and alternative sources of genome instability in shaping fungal genome evolution.

## Materials and Methods

### Fungal cultivation

Eleven *Geosmithia* species were chosen to cover three main ecology types of the genus: three specialist species associated only with vectors feeding on plant family Pinaceae (*Geosmithia* sp. RJ08m here referred as *G. abietis* nom. prov. CCF 4201, *G. magnispora* CCF 4525, *Geosmithia* sp. CCF4223 here referred as *G. pinophila* nom. prov. CCF 4223), all three known ambrosia fungi (*G. cnesini* CCF 4292*, G. eupagioceri* CCF 3754*, G. microcorthyli* CCF 3861), and 5 generalist species with broad range of vectors and plant hosts (*G. cedri* nom. prov. CCF 4200, *G. langdonii* CCF 3332, *G. multisociorum* CCF 4528, *G. obscura* CCF 3424, *G. xerotolerans* CCF 4530). These generalist species include two sister species to ambrosial *G. microcorthyli*, *G. multisociorum* and *G. cedri* nom. prov. Strains were grown on 2% malt extract agar until the sporulation occurs. Spore suspension in water was then used to inoculate Petri dishes containing the same growth medium covered by cellophane membrane to separate mycelium from the medium. Fungi were grown for three days at 25 °C, then the mycelium was scraped off the cellophane for DNA/RNA extraction.

### DNA extraction and sequencing

DNA extraction was done using a modified version (dx.doi.org/10.17504/protocols.io.5t7g6rn) of the protocol described in (Mayjonade et al., 2016). Tris-HCl 10 mM pH=8.5 was used as an elution buffer. Purity of isolated DNA was checked on Nanodrop (NanoDrop™ 2000c, ThermoFisher Scientific). The concentration was measured using Qubit 2.0 Fluorometer (Thermo Fisher Scientific, Waltham, Massachusetts, MA, USA) and the fragment length by 0.4% gel electrophoresis and GeneRuler High Range DNA Ladder (Thermo Scientific). The fungal identity was checked by PCR amplification of ITS region and fragment sequencing. The absence of bacterial contamination was verified by PCR amplification of 16S region and subsequent electrophoresis. Total amount of DNA sent for PacBio HiFi was in range from 4,400 ng to 7,540 ng. PacBio HiFi sequencing was done at Roy J. Carver Biotechnology Center, University of Illinois Urbana-Champaign.

### RNA extraction and sequencing

RNA was extracted from 3-days culture using liquid nitrogen and RNA using the Nucleospin RNA plant kit (Macherey-Nagel). The DNA digestion step was omitted from the protocol. The DNA digestion was then performed with TURBO DNA-free kit (Invitrogen). The absence of residual DNA in the samples was verified by PCR amplification of ITS region and visualization of the reaction products on the agarose gel. The purity and concentration of isolated RNA was checked similarly as above. The RNA integrity was first check by electrophoresis for the presence of two bands belonging to 28S and 18S rRNA bands. The integrity of samples having sharp bands was then verified on Bioanalyzer using RNA 6000 Pico Kit (Agilent). Samples with RIN values higher than 6.3 were sequenced. RNA sequencing was done at Genomics and Bioinformatics Core Facility, Institute of Molecular Genetics of the CAS on Illumina NextSeq 2000 platform with read length 2 × 300 bp.

### Hi-C sequencing and analysis

DNA extraction and library preparation was done following Arima High Coverage HiC Kit protocol for Animal Tissues (Arima Genomics, A160162 v01). 150 mg of fresh weight mycelium served as an input material; 10% aliquot was then used for library preparation. Library was sequenced on Illumina NextSeq platform at CGB Genomics service facility, Indiana University. Trim Galore (Babraham Institute) was used to trim adapters and FastQC (Babraham Bioinformatics, n.d.) was used to visually check quality of sequenced reads. BWA (Li, 2013) was used to map read to assembled genomes. PCR duplicates were removed using Picard MarkDuplicates and chromosome-level scaffolding was performed with YaHS (Zhou et al., 2023), which uses Hi-C contact maps to optimize scaffolding of contigs. To enable manual curation of scaffolding, Hi-C contact matrices were constructed with the Juicer pipeline (Durand et al., 2016). The contact maps were loaded into Juicebox Assembly Tools for interactive review and manual correction of scaffolds. Manually corrected assemblies were used for further analyses. Hifiasm with Hi-C integrated assembly was used as an independent tool producing similar output assembly as the previously described method.

### Genome assembly

HiFiAdapterFilt (Sim et al., 2022) was used to remove remaining adapters from PacBio HiFi raw reads and their absence was then checked using FastQC (Babraham Bioinformatics, n.d.). Genome assembly was performed with two different assemblers. First, we used Hifiasm (Cheng et al., 2021) with default settings supplemented by -l0 option as the genomes were presumed to be homozygous, then HiCanu algorithm of Canu software (Koren et al., 2017; Nurk et al., 2020) was used with default settings and genome size= 30 Mb. Assembled genomes were compared using D-GENIES v. 1.5.0 (Cabanettes and Klopp, 2018). Both softwares produced similar assemblies; however, Hifiasm led to more contiguous assemblies for some species. Assembly quality was first check by mapping of raw reads back to assembled genome using minimap2 (Li, 2021, 2018) and visually inspected in IGV v. 2.18.2 (Robinson et al., 2023, 2011; Thorvaldsdóttir et al., 2013). Assembly statistics was performed by QUAST v. 4.6.3 (Mikheenko et al., 2016). BUSCO v. 5.7.1 (Manni et al., 2021) was used to assess genome completeness by searching for conserved genes in the Hypocreales dataset (hypocreales_odb10). The known telomere sequence was manually searches by searching for the canonical telomeric repeat motif (TTAGGG) at both 5′ and 3′ ends of the chromosomes. Before genome annotation, repetitive sequences were identified using repeatmodeler v.2.0.5 (Flynn et al., 2020) and then masked with RepeatMasker http://repeatmasker.org.

### Genome annotation

Genome annotation was done using RNA seq and custom protein database as evidence. First RNA raw reads were checked using FastQC (Babraham Bioinformatics, n.d.). As most of reads contained long polyA and polyG tails we used Trim Galore (Babraham Institute) with option --hardtrim5 100 to trim read length to 100 bases. The trimmed reads were then mapped to assembled genomes using hisat2 v2.2.1 (Kim et al., 2019). Custom protein database was constructed from annotated proteins of reference strains belonging to Hypocreales available on NCBI on 10.12.2024. Soft-masked genomes were used as an input file for BRAKER3 pipeline (Gabriel et al., 2024). Final annotation was ran using the default settings was supplemented by --softmasking --gm_max_intergenic 5000 --gff3 --fungus -- busco_lineage=hypocreales_odb10 options. Annotation without busco_lineage option was run to assess the quality of annotation by checking the BUSCO completeness.

### Phylogenetics

Phylogenetic three was constructed from 3,816 single copy BUSCO genes. BUSCO Phylogenomics pipeline (https://github.com/jamiemcg/BUSCO_phylogenomics) was used to extract single copy protein sequences from BUSCO analysis outputs and generate multiple sequence alignments with MUSCLE (Edgar, 2004). These alignments were then trimmed using trimAI (Capella-Gutiérrez et al., 2009) and trimmed alignments were then concatenated into supermatrix alignment. Apart from eleven *Geosmithia* species sequenced in the present study, genome of *G. morbida* (GCF_ 012550715.1, (Aggarwal et al., 2016)) and *Emericellopsis terricola* (GCA_027569275.1, (Caesar et al., 2023)) were downloaded from NCBI and included into the analysis. Iqtree2 (Minh et al., 2020) was used to build individual gene trees and species tree from supermatrix alignment (Chernomor et al., 2016) using ultrafast bootstrap – 1000 (Hoang et al., 2018) and -alrt 1000. Best model for each gene was identified using ModelFinder (Kalyaanamoorthy et al., 2017) as a part of iqtree2 run. Iqtree2 was also used to calculate the gene and site concordance factor (Mo et al., 2023) between species and gene trees. All trees were rooted on *E. terricola*.

### Orthology structural analyses and putative centromere identification

Gene orthology was assessed by OrthoFinder v. 2.5.4 (Emms and Kelly, 2015) using the default settings. Single copy genes were then used to build and visualize chromosome synteny using GENESPACE (Lovell et al., 2022). Genome of *Geosmithia magnispora* was used as the reference and the ordering of species followed their phylogeny.

We calculated the GC% and repetitive elements and gene density along chromosomes by dividing genomes in windows of 20 kb using the utilities makewindows, nuc and coverage of BEDTools v. 2.31.0 (Quinlan, 2014). Centromere positions were inferred directly from Hi-C contact maps, as centromeres are known to produce characteristic interaction patterns marked by increased chromatin contacts at their locations. All genomic features were visualized using karyoploteR (Gel and Serra, 2017). Syntenic blocks in phylogenetic lineages were taken from GENESPACE output syntenicHits.

### Centromere characterization

Centromere positions were inferred from chromosome-scale Hi-C contact maps generated for four species: *Geosmithia eupagioceri, G. multisociorum, G. microcorthyli,* and *G. pinophila* nom.prov. In fungi, centromeric chromatin typically shows strong local self-interaction which on Hi-C maps appears as focal contact enrichments along the diagonal. Consistently with the Rabl chromosome configuration, in which centromeres and telomeres occupy opposite nuclear poles, centromeres and telomeres interact with each other on a long-distance range, which is displayed as discrete off-diagonal interaction dots at the intersections of their coordinates. Putative centromere positions were identified from these recurrent interaction signals and subsequently inspected for repeat content, gene density, transcriptional status, and synteny conservation across species.

To identify the centromeric histone, annotated proteomes were first screened for CenH3 homologs. Because no obvious CenH3 candidate was detected, additional similarity searches were performed using functionally characterized CenH3 from *Neurospora crassa* as a query in tblastn search. Additionally, we compiled reference datasets of annotated CenH3 and canonical H3 proteins from Sordariomycetes proteomes available on NCBI database. Hidden Markov Model (HMM)-based searches were then performed against annotated proteins from *Geosmithia morbida* downloaded from NCBI database and the proteomes of 11 *Geosmithia* species from the current study. Recovered sequences were aligned using MAFFT followed by phylogenetic reconstruction to distinguish CenH3 from canonical H3 lineages.

### Functional analysis

Gene functions were assigned using eggNOG-mapper v2.1.8 with the eggNOG orthology v5.0.2 database on Galaxy Europe. Carbohydrate-Active Enzymes (CAZymes) were annotated using dbCAN3, [dbCAN3 server](https://pro.unl.edu/dbCAN2/), (Zhen, g et al., 2023) with HMMER, Diamond, and dbCAN_sub. Only CAZymes genes annotated by at least two tools were included in subsequent analyses. CAZymes were divided into six classes following CAZY (Carbohydrate-Active enZymes) Database, namely Glycoside Hydrolases (GHs), GlycosylTransferases (GTs), Polysaccharide Lyases (PLs), Carbohydrate Esterases (CEs), Auxiliary Activities (AAs),and Carbohydrate-Binding Modules (CBMs), and their respective families. Genes annotated into KEGG_ko categories by eggnog-mapper or dbCAN3 were used KEGG Mapper, [KEGG PATHWAY Database](https://www.kegg.jp/kegg/pathway.html), to identify metabolic pathways in which they are involved. BlastKOALA tool, [BlastKOALA](https://www.kegg.jp/blastkoala/), was used to divide annotated proteins into the main metabolic functions.

Signal proteins were first predicted with SignalP 6.0 (Teufel et al., 2022). Signal proteins with transmembrane helices were detected with Phobius 1.01 (Käll et al., 2004) and DeepTMHMM 1.0 (Hallgren et al., 2022), and proteins with GPI anchors were identified using PredGPI (Pierleoni et al., 2008). These proteins were excluded from further analyses. Localization of remaining proteins was inferred using DeepLoc 2.1 (Ødum et al., 2024); only proteins with predicted extracellular localization were further used as an input for analysis of putative effectors using EffectorP 3.0 (Sperschneider and Dodds, 2022). A BlastP search against the PHI-base database v.4.18 (Urban et al., 2025) was used to track genes involved in pathogen-host interactions. Biosynthetic gene clusters (BGCs) were predicted using fungal version antiSMASH v. 8.0 (Blin et al., 2025), [antiSMASH fungal version](https://fungismash.secondarymetabolites.org/#!/start), with the detection strictness set to relaxed. The assembled genomes and their respective gff3 file were used as input files.

Repeat-induced point (RIP) mutation estimation

The RIP profile was assessed using the online version of the RIPper (Van Wyk et al., 2019), (https://theripper.hawk.rocks/), using default settings with minimum composite value set on 0.01, minimum product higher than 1.1, maximum substrate smaller than 0.75, and composite index chain at least 7. Sliding window approach was used with 1,000 bp windows and 500 bp step sizes. The occurrence and extend of RIP were classified based on (van Wyk et al., 2021) into six classes based on the proportion of RIP affected genome: class 1 – no RIP (0.0 to < 0.2%), Class 2 – low RIP (0.2 to <1.0%), Class 3 – moderately low RIP (1.0 to<5.0%), Class 4 – moderate RIP (5.0 to<10.0%), Class 5 – moderately high RIP (10.0 to <20.0%), and Class 6 high RIP (≥20.0%).

### Gene family evolution

We estimated gene family size expansion (gains) and contraction (losses) on each branch using CAFE5 (Mendes et al., 2020). The input files consisted of the number of genes per species belonging to each orthogroup defined by OrthoFinder and an ultrametric phylogenetic tree. Only orthogroups present in at least two *Geosmithia* species were considered. The ultrametric phylogeny of *Geosmithia* species was obtained from a maximum-likelihood species tree inferred from concatenated BUSCO single-copy orthologs and subsequently converted to an ultrametric tree in R using the chronos() function from the package ape. Branch divergence times were scaled relative to an arbitrary root age set to 100.

To incorporate among-family rate variation, several *k* values were tested. The option *-k* 2 was selected as the optimal setting as it improved the likelihood compared to *-k* 1, while higher *k* values produced inconsistent results. The *-e* option was first run without a provided file to estimate the global error model, and the resulting error model was then used in all subsequent analyses.

### Selection analyses

We used PRANK v.250331 (Löytynoja, 2013) to perform multiple codon-aware protein-coding nucleotide sequence alignments of single-copy orthologs. To guide the alignment process, a fixed species phylogeny inferred using IQ-TREE on BUSCO single copy gene dataset was supplied via the -t option. The-once flag was applied to force PRANK to use the supplied tree topology without re-estimation. The -F option was used to prevent trimming of sequence ends. Next we applied BUSTED (Branch-Site Unrestricted Statistical Test for Episodic Diversification) implemented in HyPhy v. 2.5 (Kosakovsky Pond et al., 2020) to perform a gene-wide test for evidence of positive selection across the entire phylogeny. Finally, orthogroups showing signatures of positive selection were extracted and used as an input for aBSREL (adaptive Branch-Site Random Effects Likelihood) of the same tool. We conducted an exploratory analysis in which all branches were tested for positive selection (Smith et al., 2015).

## Data access

The datasets presented in this study can be found in online repositories. The genome assemblies and raw sequencing reads have been deposited in the European Nucleotide Archive (ENA) under BioProject accession PRJEB123037. Genome assemblies and annotations, together with the corresponding BED files, predicted coding sequences (CDS), and predicted protein sequences, are publicly available on Figshare under DOI: 10.6084/m9.figshare.33455809.

## Acknowledgements

This study was supported by the project Strategie AV21 project “VP33 MycoLife – the world of fungi” of the Czech Academy of Sciences and by a research fellowship granted by the Fulbright Commission in the Czech Republic. Ryan Bracewell was supported by NIH NIGMS MIRA grant: R35GM151123. Computational resources were provided by the Indiana University UITS High Performance Computing Cluster. Sequencing was carried out at the Center for Genomics and Bioinformatics at Indiana University. We thank Matt Hahn for discussions about CAFE analysis. We thank Valerie Veselská for graphical support. Author contribution: Conceptualization: TV, RB. Data curation: TV, RB. Formal analysis: TV. Funding acquisition: TV, RB, MK. Investigation: TV. Methodology: TV, RB. Project administration: TV, RB. Supervision: TV, RB. Validation: TV, RB. Visualization: TV, RB. Writing – original draft: TV. Writing – review and editing: TV, RB, MK.

