## Supplemental Figs for "Genome reshuffling as a route to specialization? Chromosome-level insights from the bark beetle symbiont genus *Geosmithia*"

**Supplementary Materials**

**Table of content**

**Supplementary Figure 1**. Genome characteristics of 11 ***Geosmithia*** species.

**Supplementary Figure 2**. KEGG pathway reconstruction for biosynthesis of individual amino acids conserved in all *Geosmithia* species.

**Supplementary Figure 3**. KEGG pathway reconstruction for biosynthesis of vitamins conserved in all *Geosmithia* species.

**Supplementary Figure 4**: Genomic landscapes of *Geosmithia* species.

**Supplementary Figure 5**: Hi-C contact maps for four *Geosmithia* species and genomic profiles along chromosomes.

**Supplementary Table 1**. CAZyme families

**Supplementary Table 2**. Signal protein analysis.

**Supplementary Table 3.** Secondary metabolite clusters

**Supplementary Table 4.** Pathogen-host interaction analysis.

**Supplementary Table 5**. The output from OrthoFinder analysis.

**Supplementary Figure 1.** Genome characteristics of 11 ***Geosmithia*** species**.** Each plot shows genomic features across chromosomes/contigs. Tracks are arranged from the outer to inner circle as follows: numbered chromosomes/contigs, gene density, repeat density, GC content, and illustration of species. Gene density, repeat density, and GC content were calculated in 20 kb windows. Bars indicate 10 µm. Illustrations by Valerie Veselská.

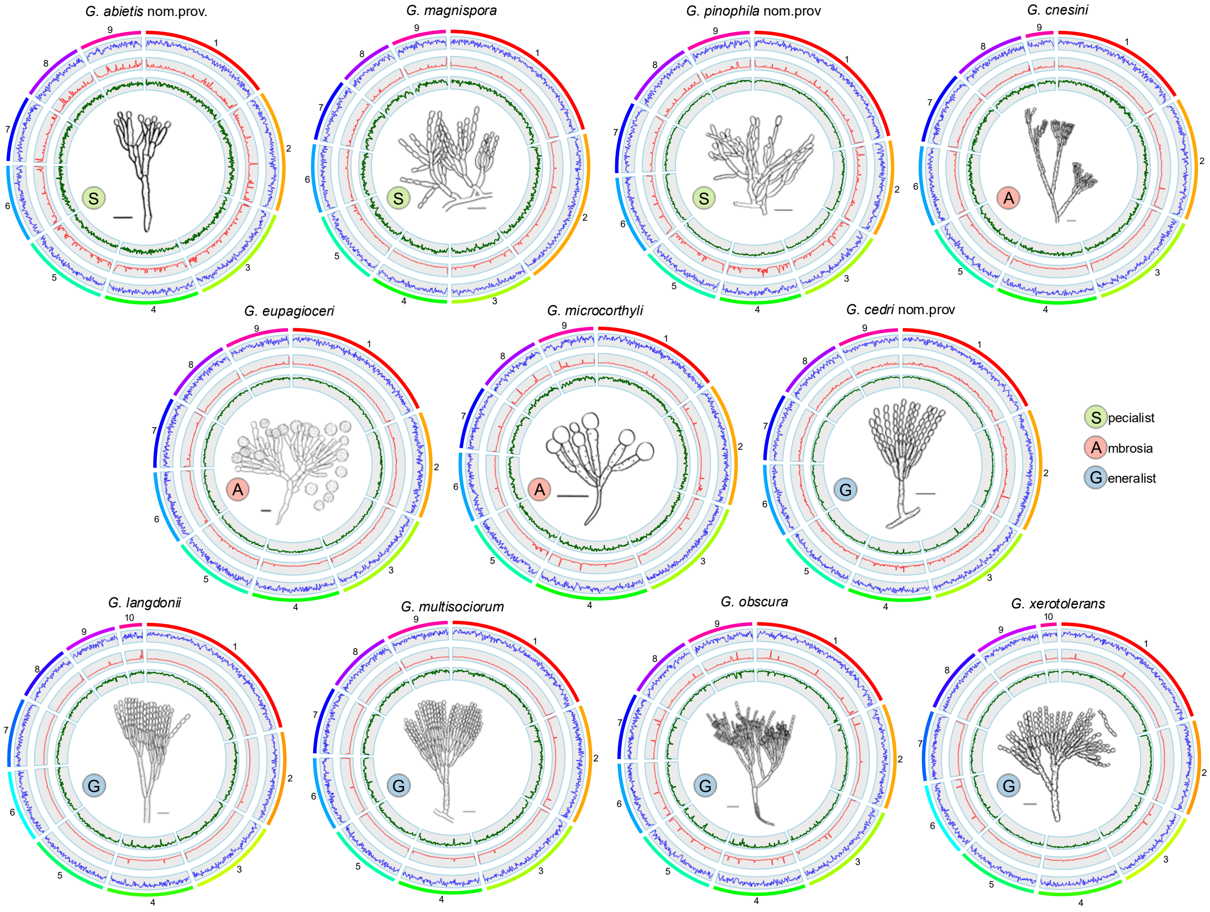

**Supplementary Figure 2**. KEGG pathway reconstruction for biosynthesis of individual amino acids conserved in all *Geosmithia* species.

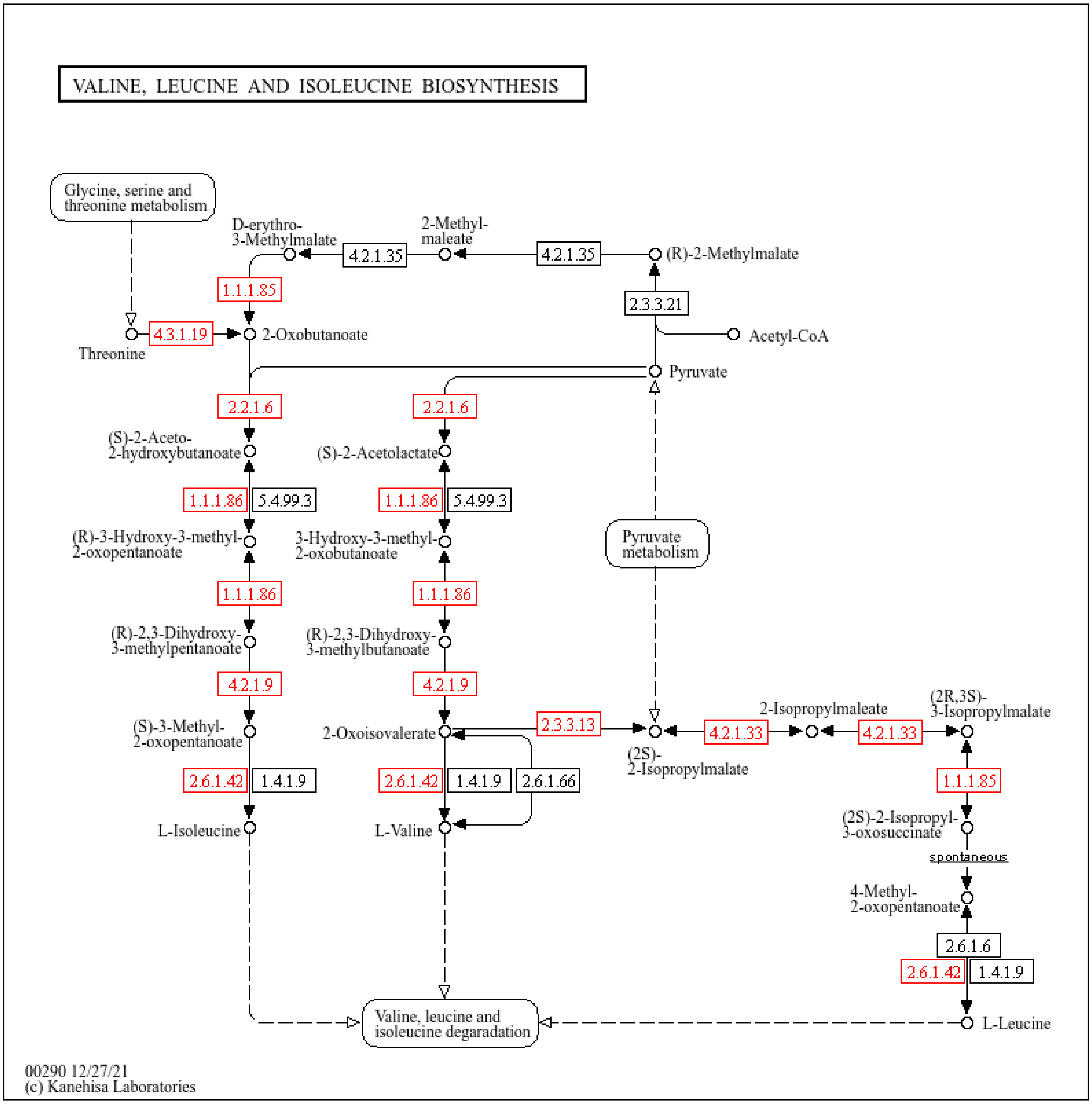

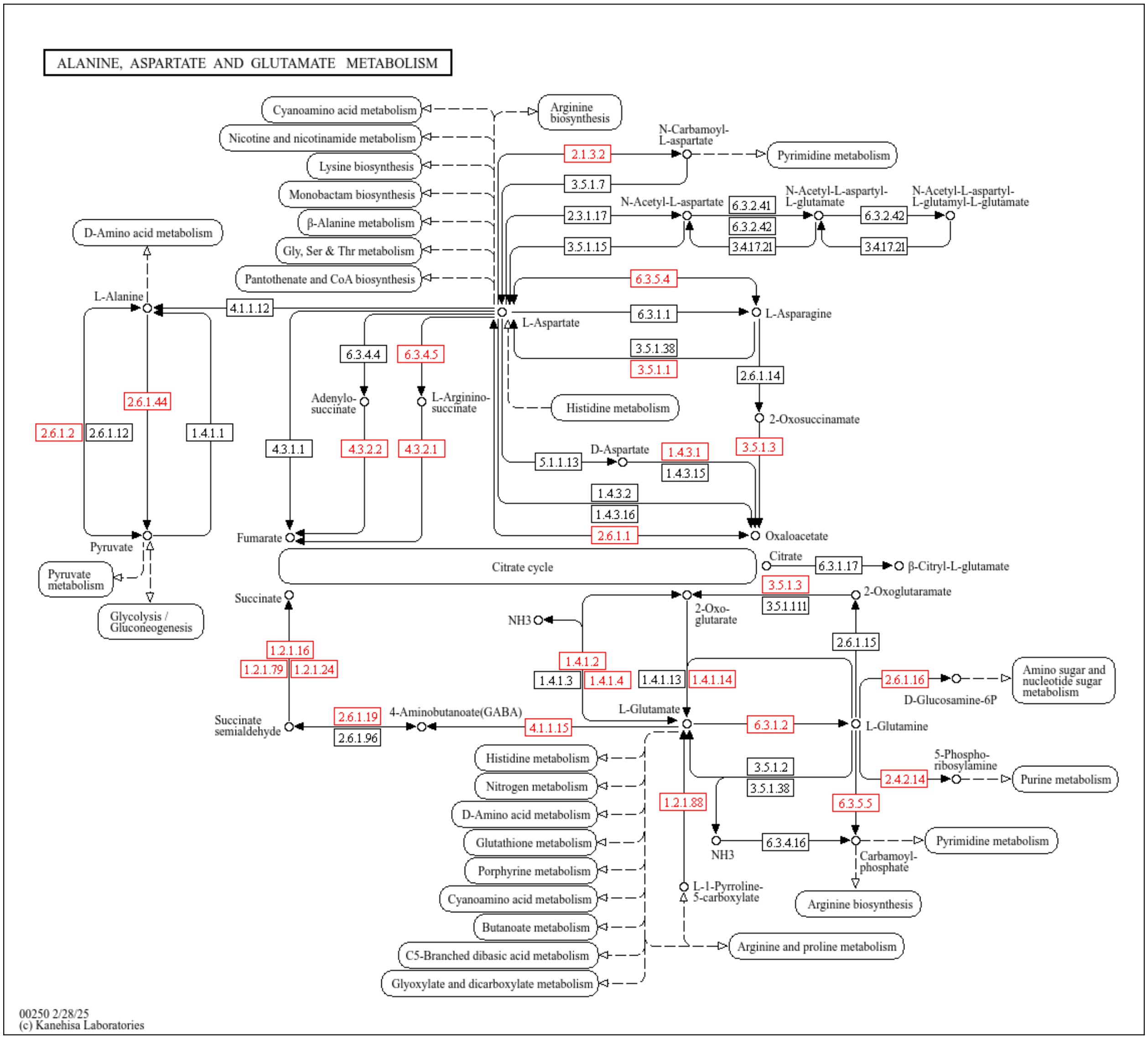
ALANINE, ASPARTATE AND GLUTAMATE METABOLISM
Cyanoamino acid metabolism
Nicotine and nicotinamide metabolism
Lysine biosynthesis
Monobactam biosynthesis
β-Alanine metabolism
Gly, Ser & Thr metabolism
Pantothenate and CoA biosynthesis
Arginine biosynthesis
N-Carbamoyl-L-aspartate
Pyrimidine metabolism
2.1.3.2
3.5.1.7
2.3.1.17
3.5.1.15
N-Acetyl-L-aspartate
6.3.2.41
6.3.2.42
3.4.17.21
N-Acetyl-L-aspartyl-L-glutamate
6.3.2.42
3.4.17.21
N-Acetyl-L-aspartyl-L-glutamyl-L-glutamate
D-Amino acid metabolism
L-Alanine
4.1.1.12
6.3.5.4
L-Aspartate
6.3.1.1
L-Asparagine
3.5.1.38
3.5.1.1
6.3.4.4
6.3.4.5
Adenylo-succinate
L-Arginino-succinate
Histidine metabolism
2.6.1.14
2-Oxosuccinamate
2.6.1.44
2.6.1.2
2.6.1.12
1.4.1.1
4.3.1.1
4.3.2.2
4.3.2.1
3.5.1.3
5.1.1.13
D-Aspartate
1.4.3.1
1.4.3.15
1.4.3.2
1.4.3.16
2.6.1.1
Pyruvate
Fumarate
Oxaloacetate
Pyruvate metabolism
Glycolysis / Gluconeogenesis
Citrate cycle
Citrate
6.3.1.17
β-Citryl-L-glutamate
3.5.1.3
3.5.1.111
2-Oxoglutaramate
Succinate
NH3
2-Oxo-glutarate
2.6.1.15
1.2.1.16
1.2.1.79
1.2.1.24
1.4.1.2
1.4.1.3
1.4.1.4
1.4.1.13
1.4.1.14
2.6.1.16
D-Glucosamine-6P
Amino sugar and nucleotide sugar metabolism
Succinate semialdehyde
2.6.1.19
2.6.1.96
4-Aminobutanoate(GABA)
4.1.1.15
L-Glutamate
6.3.1.2
L-Glutamine
5-Phospho-ribosylamine
Purine metabolism
2.4.2.14
3.5.1.2
3.5.1.38
Histidine metabolism
Nitrogen metabolism
D-Amino acid metabolism
Glutathione metabolism
Porphyrine metabolism
Cyanoamino acid metabolism
Butanoate metabolism
C5-Branched dibasic acid metabolism
Glyoxylate and dicarboxylate metabolism
1.2.1.88
6.3.5.5
NH3
6.3.4.16
Pyrimidine metabolism
Carbamoyl-phosphate
L-1-Pyrroline-5-carboxylate
Arginine biosynthesis
Arginine and proline metabolism
00250 2/28/25
(c) Kanehisa Laboratories

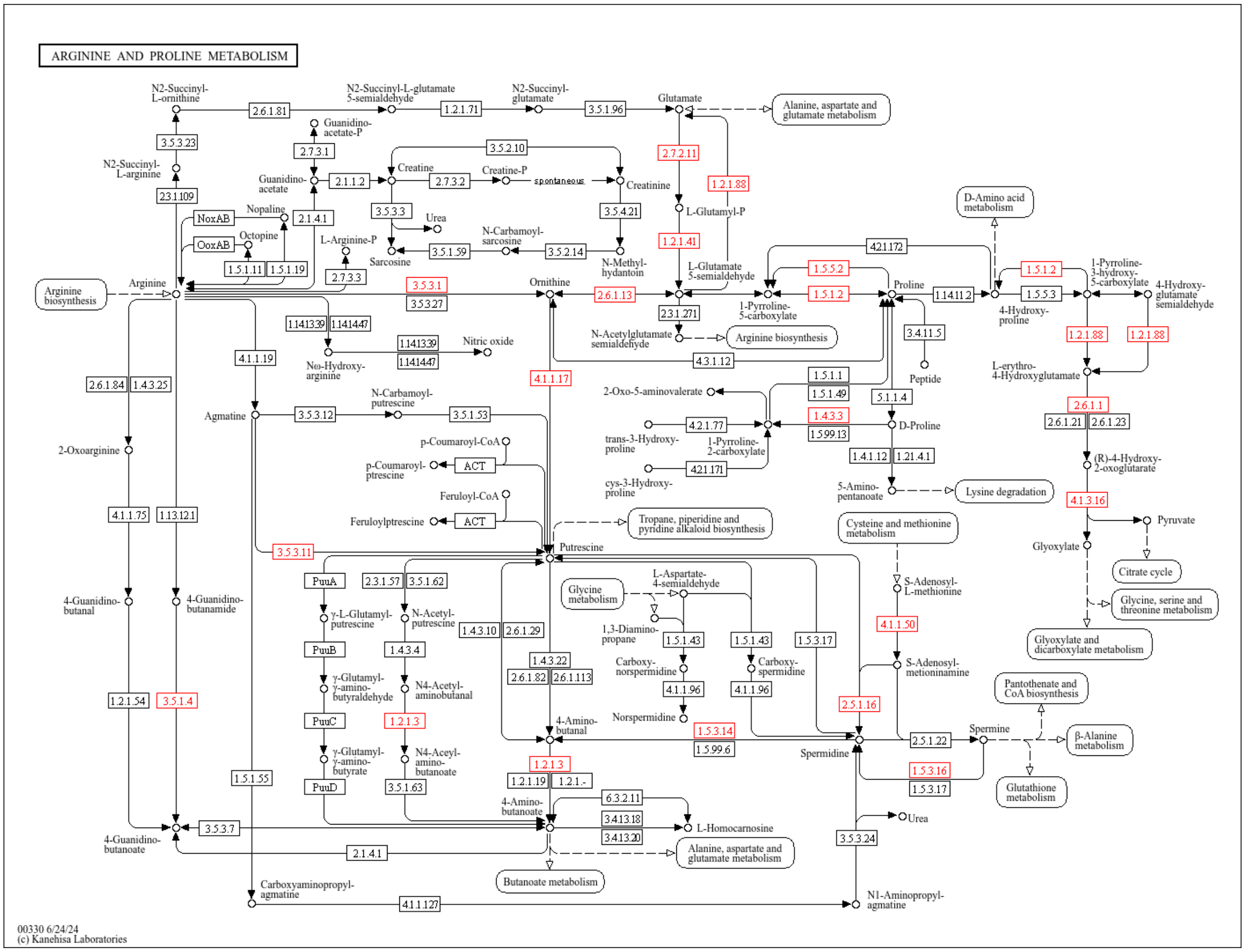

ARGININE AND PROLINE METABOLISM
N2-Succinyl-L-ornithine
2.6.1.81
N2-Succinyl-L-glutamate 5-semialdehyde
1.2.1.71
N2-Succinyl-glutamate
3.5.1.96
Glutamate
Alanine, aspartate and glutamate metabolism
3.5.3.23
Guanidino-acetate-P
2.7.3.1
3.5.2.10
N2-Succinyl-L-arginine
Creatine
Creatine-P
2.7.2.11
Guanidino-acetate
2.1.1.2
2.7.3.2
spontaneous
Creatinine
1.2.1.88
2.3.1.109
L-Glutamyl-P
NoxAB
Nopaline
2.1.4.1
3.5.3.3
3.5.4.21
Urea
D-Amino acid metabolism
OoxAB
Octopine
L-Arginine-P
N-Carbamoyl-sarcosine
1.2.1.41
4.2.1.172
1.5.1.11
1.5.1.19
3.5.1.59
3.5.2.14
Sarcosine
N-Methyl-hydantoin
L-Glutamate 5-semialdehyde
1.5.5.2
1.5.1.2
1-Pyrroline-3-hydroxy-5-carboxylate
2.7.3.3
4-Hydroxy-glutamate semialdehyde
Arginine biosynthesis
Arginine
3.5.3.1
Ornithine
2.6.1.13
1.5.1.2
Proline
1.14.11.2
1.5.5.3
3.5.3.27
2.3.1.271
1-Pyrroline-5-carboxylate
4-Hydroxy-proline
1.14.13.39
1.14.14.47
N-Acetylglutamate semialdehyde
Arginine biosynthesis
3.4.11.5
1.2.1.88
1.2.1.88
1.14.13.39
Nitric oxide
1.14.14.47
4.1.1.19
Nω-Hydroxy-arginine
4.3.1.12
Peptide
L-erythro-4-Hydroxyglutamate
4.1.1.17
1.5.1.1
2.6.1.84
1.4.3.25
2-Oxo-5-aminovalerate
1.5.1.49
5.1.1.4
2.6.1.1
N-Carbamoyl-putrescine
Agmatine
3.5.3.12
3.5.1.53
1.4.3.3
D-Proline
2.6.1.21
2.6.1.23
4.2.1.77
1.5.99.13
p-Coumaroyl-CoA
trans-3-Hydroxy-proline
1-Pyrroline-2-carboxylate
2-Oxoarginine
1.4.1.12
1.2.1.41
(R)-4-Hydroxy-2-oxoglutarate
p-Coumaroyl-ptrescine
ACT
4.2.1.171
cys-3-Hydroxy-proline
Feruloyl-CoA
5-Amino-pentanoate
Lysine degradation
4.1.3.16
4.1.1.75
1.13.12.1
Feruloylptrescine
ACT
Tropane, piperidine and pyridine alkaloid biosynthesis
Cysteine and methionine metabolism
Pyruvate
Glyoxylate
3.5.3.11
Putrescine
Citrate cycle
PuuA
2.3.1.57
3.5.1.62
L-Aspartate-4-semialdehyde
S-Adenosyl-L-methionine
4-Guanidino-butanal
4-Guanidino-butanamide
Glycine metabolism
Glycine, serine and threonine metabolism
γ-L-Glutamyl-putrescine
N-Acetyl-putrescine
1.4.3.10
2.6.1.29
1,3-Diamino-propane
4.1.1.50
PuuB
1.4.3.4
1.5.1.43
1.5.1.43
1.5.3.17
Glyoxylate and dicarboxylate metabolism
1.4.3.22
Carboxy-norspermidine
Carboxy-spermidine
S-Adenosyl-methioninamine
γ-Glutamyl-γ-amino-butyraldehyde
N4-Acetyl-aminobutanal
2.6.1.82
2.6.1.113
4.1.1.96
4.1.1.96
Pantothenate and CoA biosynthesis
1.2.1.54
3.5.1.4
2.5.1.16
PuuC
1.2.1.3
4-Amino-butanal
Norspermidine
1.5.3.14
Spermine
1.5.99.6
2.5.1.22
β-Alanine metabolism
γ-Glutamyl-γ-amino-butyrate
N4-Acetyl-amino-butanoate
Spermidine
1.5.1.55
1.2.1.3
1.2.1.19
1.2.1.-
1.5.3.16
PuuD
3.5.1.63
1.5.3.17
Glutathione metabolism
4-Amino-butanoate
6.3.2.11
3.4.13.18
Urea
4-Guanidino-butanoate
3.5.3.7
L-Homocarnosine
3.4.13.20
3.5.3.24
2.1.4.1
Alanine, aspartate and glutamate metabolism
Butanoate metabolism
Carboxyaminopropyl-agmatine
4.1.1.127
N1-Aminopropyl-agmatine
00330 6/24/24
(c) Kanehisa Laboratories

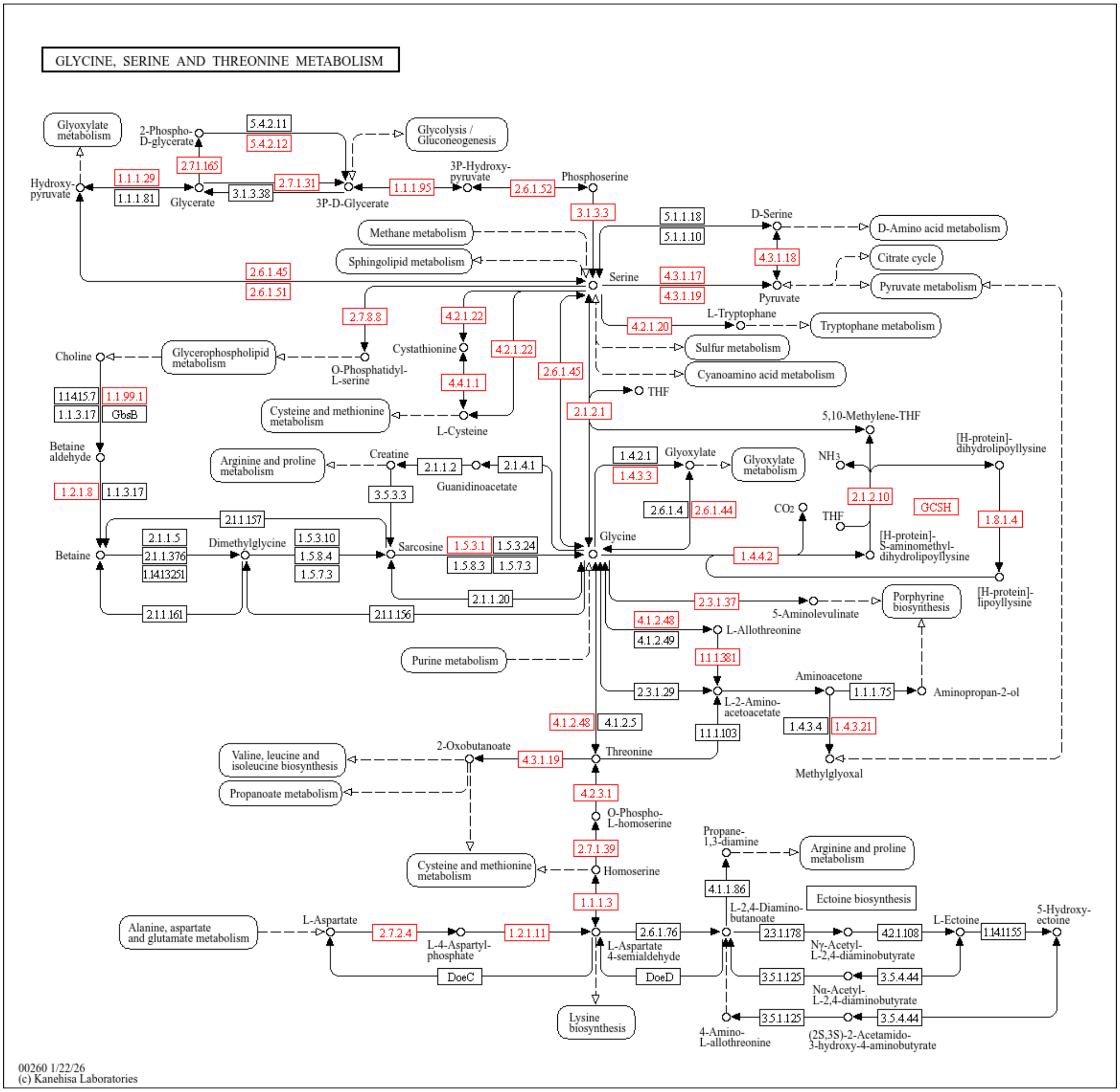
GLYCINE, SERINE AND THREONINE METABOLISM
Glyoxylate metabolism
2-Phospho-D-glycerate
5.4.2.11
5.4.2.12
Glycolysis / Gluconeogenesis
2.7.1.165
1.1.1.29
1.1.1.81
Hydroxy-pyruvate
Glycerate
3.1.3.38
2.7.1.31
3P-D-Glycerate
1.1.1.95
3P-Hydroxy-pyruvate
2.6.1.52
Phosphoserine
3.1.3.3
5.1.1.18
5.1.1.10
D-Serine
D-Amino acid metabolism
Methane metabolism
Sphingolipid metabolism
4.3.1.18
Citrate cycle
2.6.1.45
2.6.1.51
Serine
4.3.1.17
4.3.1.19
Pyruvate
Pyruvate metabolism
2.7.8.8
4.2.1.22
4.2.1.20
L-Tryptophan
Tryptophan metabolism
Sulfur metabolism
Cyanoamino acid metabolism
Choline
Glycerophospholipid metabolism
O-Phosphatidyl-L-serine
Cystathionine
4.2.1.22
2.6.1.45
4.4.1.1
THF
Cysteine and methionine metabolism
L-Cysteine
2.1.2.1
5,10-Methylene-THF
1.1.415.7
1.1.99.1
1.1.3.17
GbsB
Betaine aldehyde
Arginine and proline metabolism
Creatine
2.1.1.2
2.1.4.1
Guanidinoacetate
3.5.3.3
1.4.2.1
1.4.3.3
Glyoxylate
Glyoxylate metabolism
NH3
[H-protein]-dihydrolipoyllysine
2.1.2.10
GCSH
2.6.1.4
2.6.1.44
CO2
THF
1.8.1.4
1.2.1.8
1.1.3.17
2.1.1.157
[H-protein]-S-aminomethyl-dihydrolipoyllysine
2.1.1.5
2.1.1.376
1.14.13.251
Dimethylglycine
1.5.3.10
1.5.8.4
1.5.7.3
Sarcosine
1.5.3.1
1.5.3.24
1.5.8.3
1.5.7.3
Glycine
1.4.4.2
Betaine
2.1.1.20
[H-protein]-lipoyllysine
2.3.1.37
5-Aminolevulinate
Porphyrine biosynthesis
2.1.1.161
2.1.1.156
4.1.2.48
4.1.2.49
L-Allothreonine
Purine metabolism
1.1.1.381
Aminoacetone
2.3.1.29
1.1.1.75
Aminopropan-2-ol
L-2-Amino-acetoacetate
4.1.2.48
4.1.2.5
1.1.1.103
1.4.3.4
1.4.3.21
2-Oxobutanoate
Valine, leucine and isoleucine biosynthesis
4.3.1.19
Threonine
Methylglyoxal
Propanoate metabolism
4.2.3.1
O-Phospho-L-homoserine
2.7.1.39
Propane-1,3-diamine
Arginine and proline metabolism
Cysteine and methionine metabolism
Homoserine
4.1.1.86
Ectoine biosynthesis
1.1.1.3
L-2,4-Diamino-butanoate
5-Hydroxy-ectoine
Alanine, aspartate and glutamate metabolism
L-Aspartate
2.7.2.4
1.2.1.11
2.6.1.76
2.3.1.178
4.2.1.108
L-Ectoine
1.14.11.55
L-4-Aspartyl-phosphate
L-Aspartate 4-semialdehyde
Nγ-Acetyl-L-2,4-diaminobutyrate
DoeC
DoeD
3.5.1.125
3.5.4.44
Nα-Acetyl-L-2,4-diaminobutyrate
Lysine biosynthesis
3.5.1.125
3.5.4.44
4-Amino-L-allothreonine
(2S,3S)-2-Acetamido-3-hydroxy-4-aminobutyrate
00260 1/22/26
(c) Kanehisa Laboratories

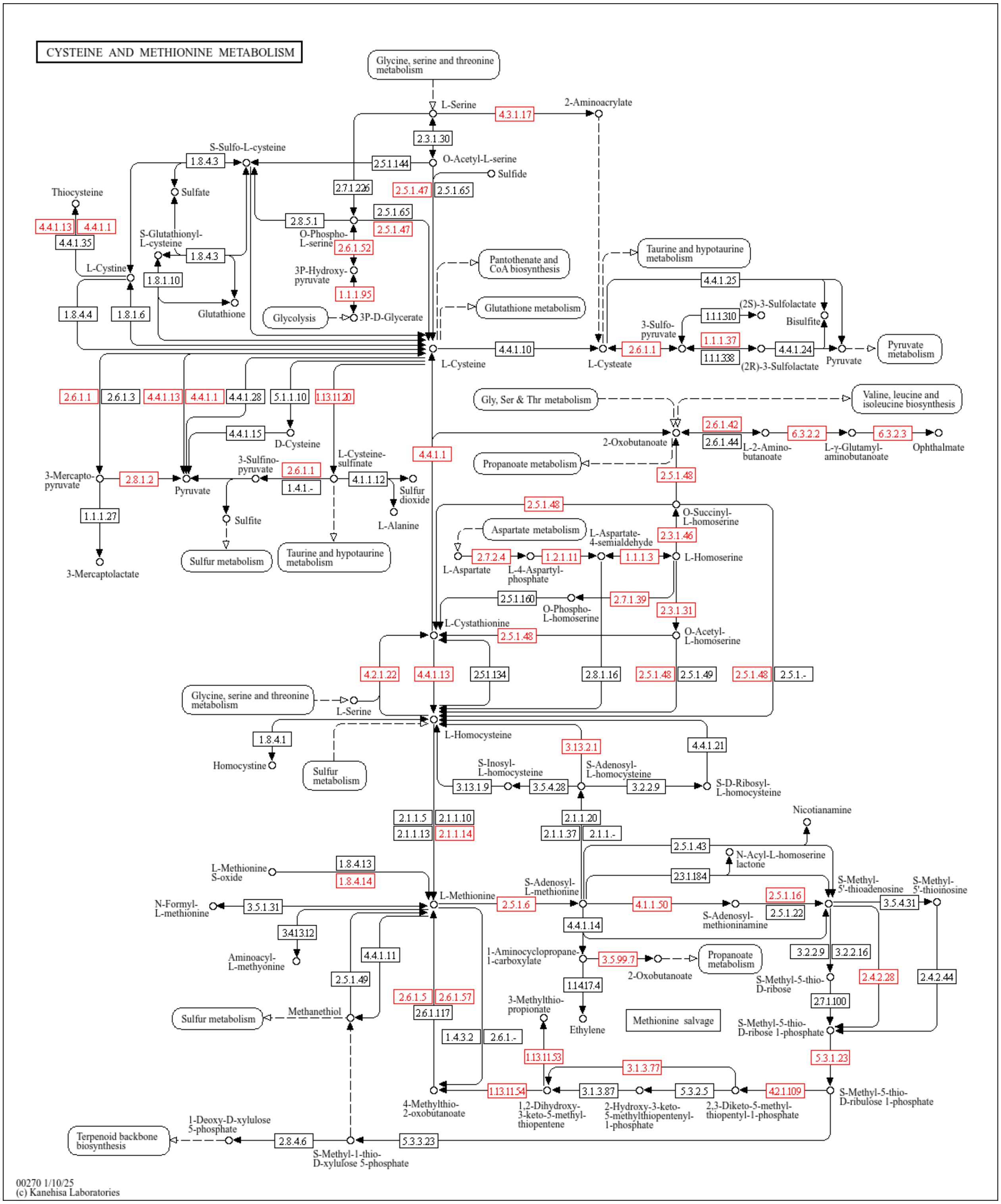
CYSTEINE AND METHIONINE METABOLISM
Glycine, serine and threonine metabolism
L-Serine
4.3.1.17
2-Aminoacrylate
2.3.1.30
S-Sulfo-L-cysteine
1.8.4.3
2.5.1.144
O-Acetyl-L-serine
Sulfide
Thiocysteine
Sulfate
2.7.1.226
2.5.1.47
2.5.1.65
4.4.1.13
4.4.1.1
2.8.5.1
2.5.1.65
4.4.1.35
S-Glutathionyl-L-cysteine
1.8.4.3
O-Phospho-L-serine
2.5.1.47
2.6.1.52
Pantothenate and CoA biosynthesis
Taurine and hypotaurine metabolism
L-Cystine
1.8.1.10
3P-Hydroxy-pyruvate
4.4.1.25
1.1.1.95
Glutathione metabolism
(2S)-3-Sulfolactate
1.1.1.310
Bisulfite
1.8.4.4
1.8.1.6
Glutathione
Glycolysis
3P-D-Glycerate
3-Sulfo-pyruvate
1.1.1.37
4.4.1.10
2.6.1.1
4.4.1.24
Pyruvate metabolism
1.1.1.338
L-Cysteine
L-Cysteate
(2R)-3-Sulfolactate
Pyruvate
2.6.1.1
2.6.1.3
4.4.1.13
4.4.1.1
4.4.1.28
5.1.1.10
1.13.11.20
Gly, Ser & Thr metabolism
Valine, leucine and isoleucine biosynthesis
4.4.1.15
D-Cysteine
2.6.1.42
6.3.2.2
6.3.2.3
2-Oxobutanoate
2.6.1.44
L-2-Amino-butanoate
L-γ-Glutamyl-aminobutanoate
Ophthalmate
4.4.1.1
L-Cysteine-sulfinate
3-Sulfino-pyruvate
Propanoate metabolism
3-Mercapto-pyruvate
2.8.1.2
Pyruvate
2.6.1.1
1.4.1.-
4.1.1.12
Sulfur dioxide
2.5.1.48
1.1.1.27
Sulfite
L-Alanine
2.5.1.48
O-Succinyl-L-homoserine
Aspartate metabolism
L-Aspartate-4-semialdehyde
2.3.1.46
3-Mercaptolactate
Sulfur metabolism
Taurine and hypotaurine metabolism
2.7.2.4
1.2.1.11
1.1.1.3
L-Homoserine
L-Aspartate
L-4-Aspartyl-phosphate
2.5.1.160
2.7.1.39
O-Phospho-L-homoserine
2.3.1.31
L-Cystathionine
2.5.1.48
O-Acetyl-L-homoserine
4.2.1.22
4.4.1.13
2.5.1.134
2.8.1.16
2.5.1.48
2.5.1.49
2.5.1.48
2.5.1.-
Glycine, serine and threonine metabolism
L-Serine
L-Homocysteine
1.8.4.1
3.13.2.1
4.4.1.21
Homocystine
Sulfur metabolism
S-Inosyl-L-homocysteine
S-Adenosyl-L-homocysteine
3.13.1.9
3.5.4.28
3.2.2.9
S-D-Ribosyl-L-homocysteine
2.1.1.5
2.1.1.10
2.1.1.20
2.1.1.13
2.1.1.14
2.1.1.37
2.1.1.-
Nicotianamine
2.5.1.43
N-Acyl-L-homoserine lactone
L-Methionine S-oxide
1.8.4.13
2.3.1.184
1.8.4.14
S-Adenosyl-L-methionine
S-Methyl-5'-thioadenosine
S-Methyl-5'-thioinosine
L-Methionine
2.5.1.6
4.1.1.50
2.5.1.16
N-Formyl-L-methionine
3.5.1.31
3.5.4.31
S-Adenosyl-methioninamine
2.5.1.22
3.4.13.12
4.4.1.14
Aminoacyl-L-methyonine
4.4.1.11
1-Aminocyclopropane-1-carboxylate
3.5.99.7
Propanoate metabolism
3.2.2.9
3.2.2.16
2-Oxobutanoate
2.5.1.49
S-Methyl-5-thio-D-ribose
2.4.2.28
2.4.2.44
1.14.17.4
3-Methylthio-propionate
2.6.1.5
2.6.1.57
2.7.1.100
Methanethiol
Methionine salvage
Sulfur metabolism
2.6.1.117
Ethylene
S-Methyl-5-thio-D-ribose 1-phosphate
1.4.3.2
2.6.1.-
1.13.11.53
5.3.1.23
3.1.3.77
4-Methylthio-2-oxobutanoate
1.13.11.54
3.1.3.87
5.3.2.5
4.2.1.109
S-Methyl-5-thio-D-ribulose 1-phosphate
1,2-Dihydroxy-3-keto-5-methyl-thiopentene
2-Hydroxy-3-keto-5-methylthiopentenyl-1-phosphate
2,3-Diketo-5-methyl-thiopentyl-1-phosphate
1-Deoxy-D-xylulose 5-phosphate
Terpenoid backbone biosynthesis
2.8.4.6
5.3.3.23
S-Methyl-1-thio-D-xylulose 5-phosphate
00270 1/10/25
(c) Kanehisa Laboratories

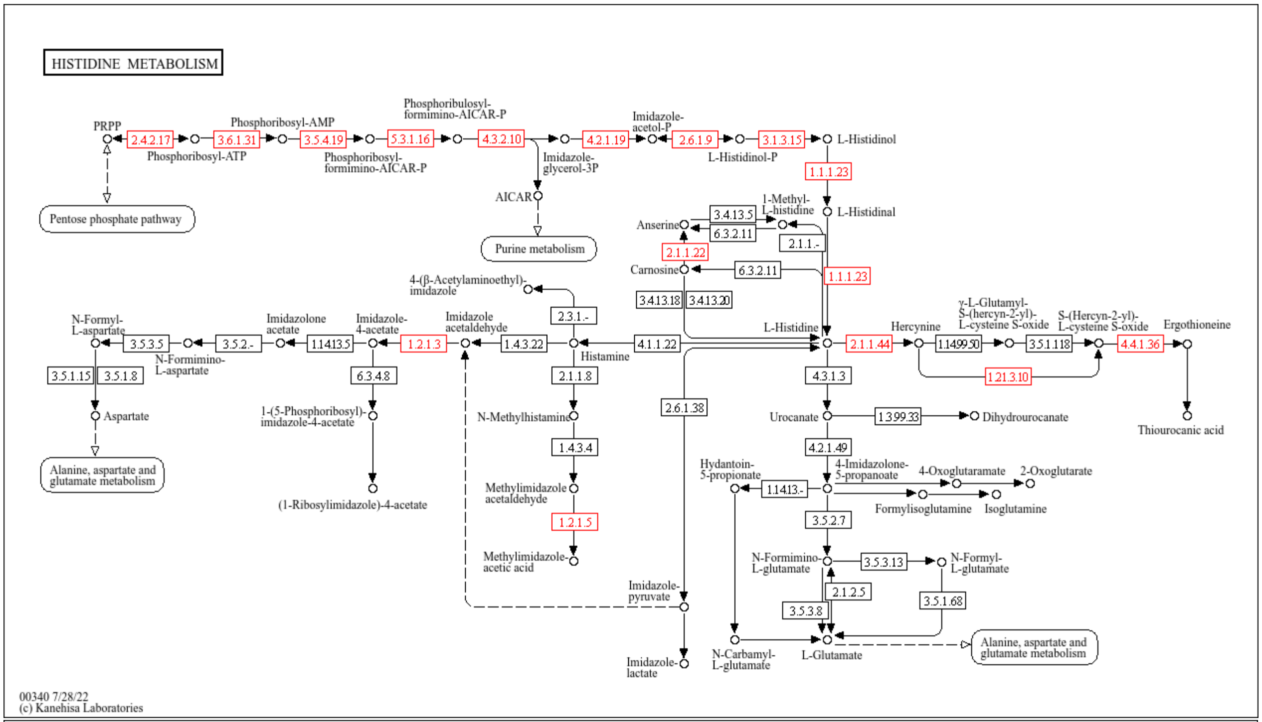
HISTIDINE METABOLISM
PRPP
2.4.2.17
Phosphoribosyl-ATP
3.6.1.31
Phosphoribosyl-AMP
3.5.4.19
Phosphoribosyl-formimino-AICAR-P
5.3.1.16
Phosphoribulosyl-formimino-AICAR-P
4.3.2.10
Imidazole-glycerol-3P
4.2.1.19
Imidazole-acetol-P
2.6.1.9
L-Histidinol-P
3.1.3.15
L-Histidinol
1.1.1.23
L-Histidinal
AICAR
Pentose phosphate pathway
Purine metabolism
Anserine
3.4.13.5
6.3.2.11
1-Methyl-L-histidine
2.1.1.-
2.1.1.22
Carnosine
6.3.2.11
1.1.1.23
3.4.13.18
3.4.13.20
4-(β-Acetylaminoethyl)-imidazole
2.3.1.-
N-Formyl-L-aspartate
3.5.3.5
N-Formimino-L-aspartate
3.5.2.-
Imidazolone acetate
1.14.13.5
Imidazole-4-acetate
1.2.1.3
Imidazole acetaldehyde
1.4.3.22
Histamine
4.1.1.22
L-Histidine
2.1.1.44
Hercynine
1.14.99.50
γ-L-Glutamyl-S-(hercyn-2-yl)-L-cysteine S-oxide
3.5.1.118
S-(Hercyn-2-yl)-L-cysteine S-oxide
4.4.1.36
Ergothioneine
1.21.3.10
Thiourocanic acid
3.5.1.15
3.5.1.8
Aspartate
6.3.4.8
1-(5-Phosphoribosyl)-imidazole-4-acetate
(1-Ribosylimidazole)-4-acetate
2.1.1.8
N-Methylhistamine
1.4.3.4
Methylimidazole acetaldehyde
1.2.1.5
Methylimidazole-acetic acid
2.6.1.38
4.3.1.3
Urocanate
1.3.99.33
Dihydrourocanate
4.2.1.49
Hydantoin-5-propionate
1.14.13.-
4-Imidazolone-5-propanoate
4-Oxoglutaramate
2-Oxoglutarate
Formylisoglutamine
Isoglutamine
3.5.2.7
N-Formimino-L-glutamate
3.5.3.13
N-Formyl-L-glutamate
2.1.2.5
3.5.3.8
3.5.1.68
Imidazole-pyruvate
N-Carbamyl-L-glutamate
L-Glutamate
Alanine, aspartate and glutamate metabolism
Alanine, aspartate and glutamate metabolism
Imidazole-lactate
00340 7/28/22
(c) Kanehisa Laboratories

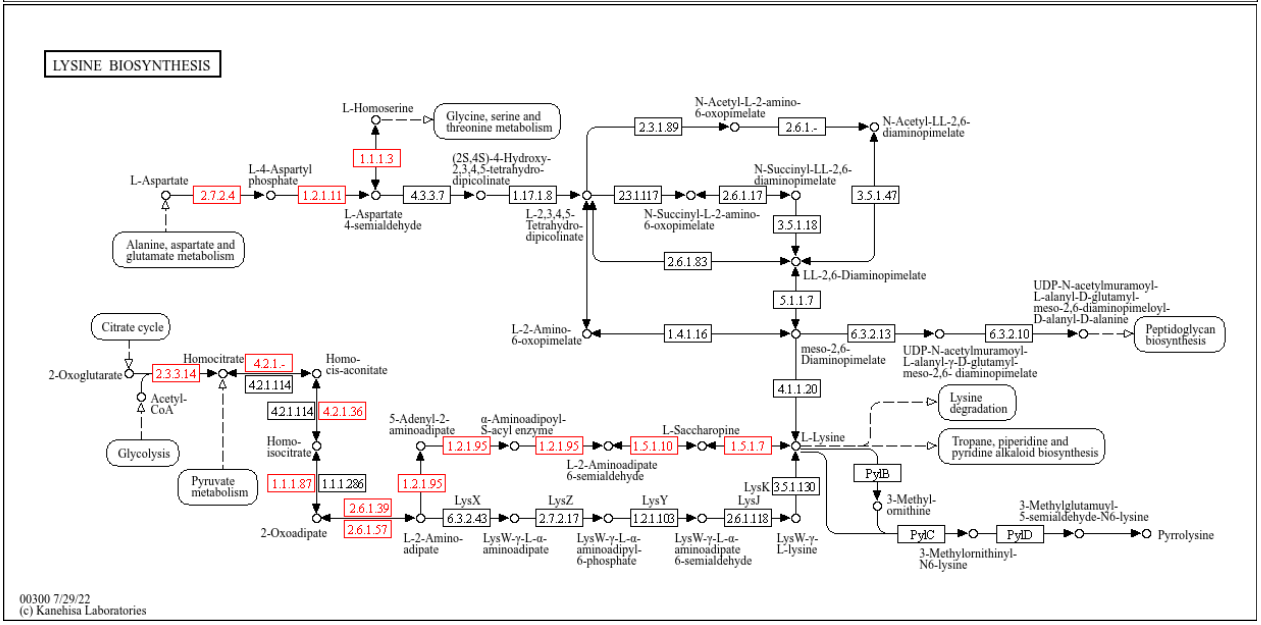
LYSINE BIOSYNTHESIS
L-Homoserine
Glycine, serine and threonine metabolism
1.1.1.3
L-Aspartate
2.7.2.4
L-4-Aspartyl phosphate
1.2.1.11
L-Aspartate 4-semialdehyde
4.3.3.7
(2S,4S)-4-Hydroxy-2,3,4,5-tetrahydro-dipicolinate
1.17.1.8
L-2,3,4,5-Tetrahydro-dipicolinate
2.3.1.89
N-Acetyl-L-2-amino-6-oxopimelate
2.6.1.-
N-Acetyl-LL-2,6-diaminopimelate
2.3.1.117
N-Succinyl-L-2-amino-6-oxopimelate
2.6.1.17
N-Succinyl-LL-2,6-diaminopimelate
3.5.1.18
3.5.1.47
2.6.1.83
LL-2,6-Diaminopimelate
Alanine, aspartate and glutamate metabolism
5.1.1.7
Citrate cycle
L-2-Amino-6-oxopimelate
1.4.1.16
meso-2,6-Diaminopimelate
6.3.2.13
UDP-N-acetylmuramoyl-L-alanyl-γ-D-glutamyl-meso-2,6- diaminopimelate
6.3.2.10
UDP-N-acetylmuramoyl-L-alanyl-D-glutamyl-meso-2,6-diaminopimeloyl-D-alanyl-D-alanine
Peptidoglycan biosynthesis
2-Oxoglutarate
2.3.3.14
Homocitrate
4.2.1.-
Homo-cis-aconitate
4.2.1.114
Acetyl-CoA
4.2.1.114
4.2.1.36
4.1.1.20
Lysine degradation
Glycolysis
Homo-isocitrate
5-Adenyl-2-aminoadipate
1.2.1.95
α-Aminoadipoyl-S-acyl enzyme
1.2.1.95
L-2-Aminoadipate 6-semialdehyde
1.5.1.10
L-Saccharopine
1.5.1.7
L-Lysine
Tropane, piperidine and pyridine alkaloid biosynthesis
Pyruvate metabolism
1.1.1.87
1.1.1.286
1.2.1.95
PylB
LysK
3.5.1.130
3-Methyl-ornithine
2.6.1.39
2.6.1.57
2-Oxoadipate
LysX
6.3.2.43
L-2-Amino-adipate
LysZ
2.7.2.17
LysW-γ-L-α-aminoadipate
LysY
1.2.1.103
LysW-γ-L-α-aminoadipyl-6-phosphate
LysJ
2.6.1.118
LysW-γ-L-α-aminoadipate 6-semialdehyde
LysW-γ-L-lysine
PylC
3-Methylornithinyl-N6-lysine
PylD
3-Methylglutamuyl-5-semialdehyde-N6-lysine
Pyrrolysine
00300 7/29/22
(c) Kanehisa Laboratories

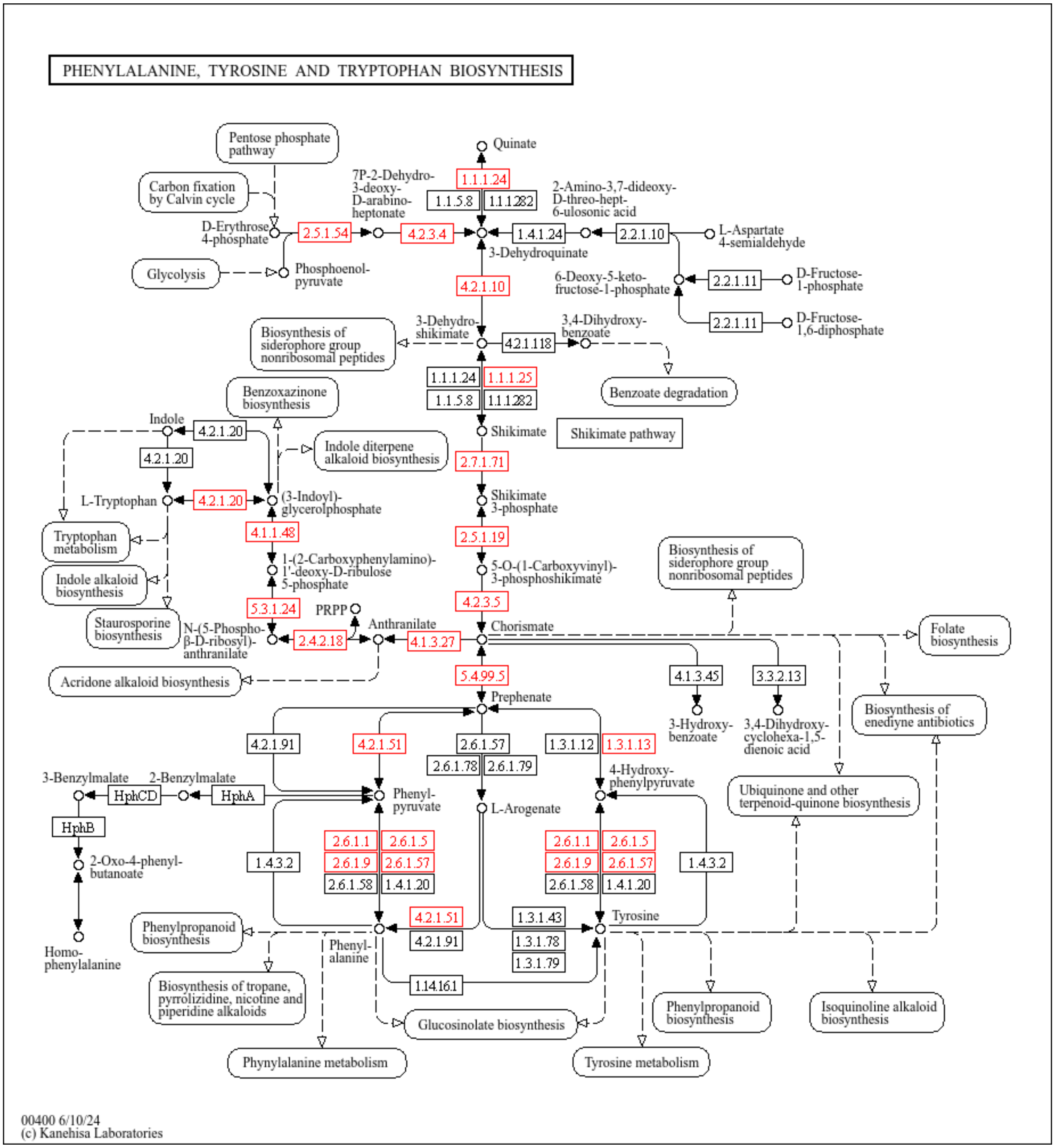
PHENYLALANINE, TYROSINE AND TRYPTOPHAN BIOSYNTHESIS
Pentose phosphate pathway
Carbon fixation by Calvin cycle
Glycolysis
D-Erythrose 4-phosphate
Phosphoenol-pyruvate
2.5.1.54
7P-2-Dehydro-3-deoxy-D-arabino-heptonate
4.2.3.4
Quinate
1.1.1.24
1.1.5.8
1.1.1.282
3-Dehydroquinate
1.4.1.24
2.2.1.10
2-Amino-3,7-dideoxy-D-threo-hept-6-ulosonic acid
L-Aspartate 4-semialdehyde
6-Deoxy-5-keto-fructose-1-phosphate
2.2.1.11
D-Fructose-1-phosphate
2.2.1.11
D-Fructose-1,6-diphosphate
4.2.1.10
3-Dehydro-shikimate
4.2.1.118
3,4-Dihydroxy-benzoate
Benzoate degradation
Biosynthesis of siderophore group nonribosomal peptides
1.1.1.24
1.1.1.25
1.1.5.8
1.1.1.282
Shikimate
Shikimate pathway
2.7.1.71
Shikimate 3-phosphate
2.5.1.19
5-O-(1-Carboxyvinyl)-3-phosphoshikimate
4.2.3.5
Chorismate
Benzoxazinone biosynthesis
Indole diterpene alkaloid biosynthesis
Indole
4.2.1.20
4.2.1.20
L-Tryptophan
4.2.1.20
(3-Indoyl)-glycerolphosphate
4.1.1.48
1-(2-Carboxyphenylamino)-1'-deoxy-D-ribulose 5-phosphate
5.3.1.24
PRPP
Tryptophan metabolism
Indole alkaloid biosynthesis
Staurosporine biosynthesis
N-(5-Phospho-β-D-ribosyl)-anthranilate
2.4.2.18
Anthranilate
4.1.3.27
Acridone alkaloid biosynthesis
Biosynthesis of siderophore group nonribosomal peptides
Folate biosynthesis
5.4.99.5
Prephenate
4.1.3.45
3.3.2.13
3-Hydroxy-benzoate
3,4-Dihydroxy-cyclohexa-1,5-dienoic acid
Biosynthesis of enediyne antibiotics
4.2.1.91
4.2.1.51
2.6.1.57
2.6.1.78
2.6.1.79
1.3.1.12
1.3.1.13
4-Hydroxy-phenylpyruvate
Ubiquinone and other terpenoid-quinone biosynthesis
3-Benzylmalate
2-Benzylmalate
HphCD
HphA
HphB
Phenyl-pyruvate
L-Arogenate
2-Oxo-4-phenyl-butanoate
1.4.3.2
2.6.1.1
2.6.1.5
2.6.1.9
2.6.1.57
2.6.1.58
1.4.1.20
2.6.1.1
2.6.1.5
2.6.1.9
2.6.1.57
2.6.1.58
1.4.1.20
1.4.3.2
Homo-phenylalanine
Phenylpropanoid biosynthesis
4.2.1.51
4.2.1.91
Phenyl-alanine
1.3.1.43
1.3.1.78
1.3.1.79
Tyrosine
1.14.16.1
Biosynthesis of tropane, pyrrolizidine, nicotine and piperidine alkaloids
Glucosinolate biosynthesis
Phenylpropanoid biosynthesis
Isoquinoline alkaloid biosynthesis
Phynylalanine metabolism
Tyrosine metabolism
00400 6/10/24
(c) Kanehisa Laboratories

**Supplementary Figure 3**. KEGG pathway reconstruction for biosynthesis of vitamins conserved in all *Geosmithia* species.

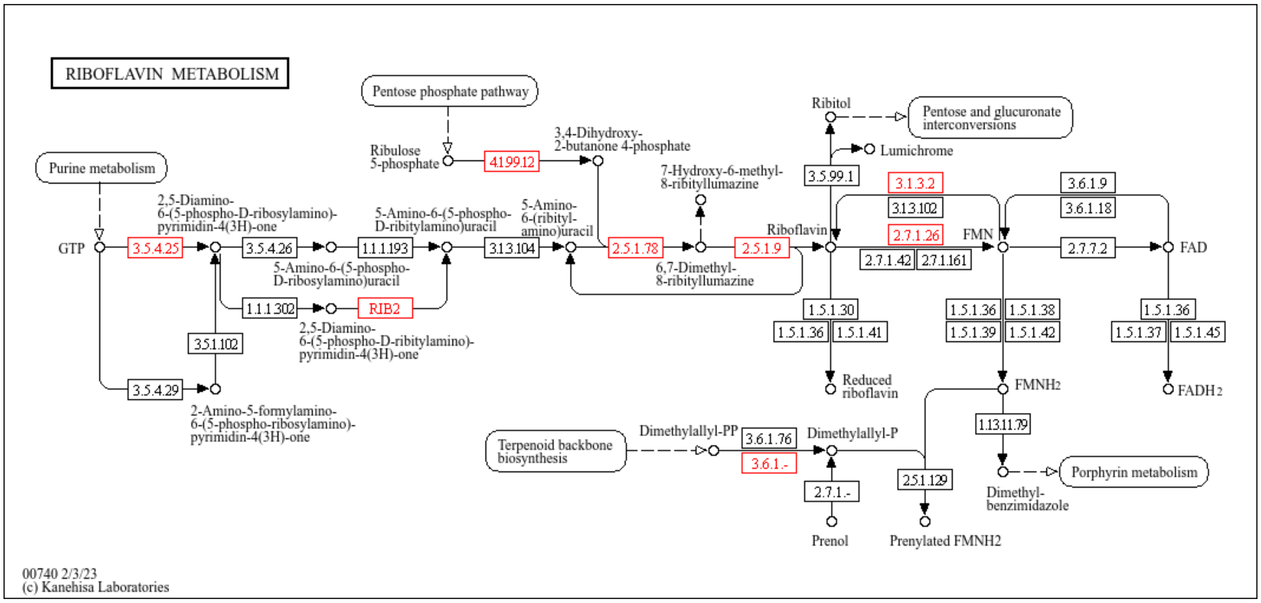


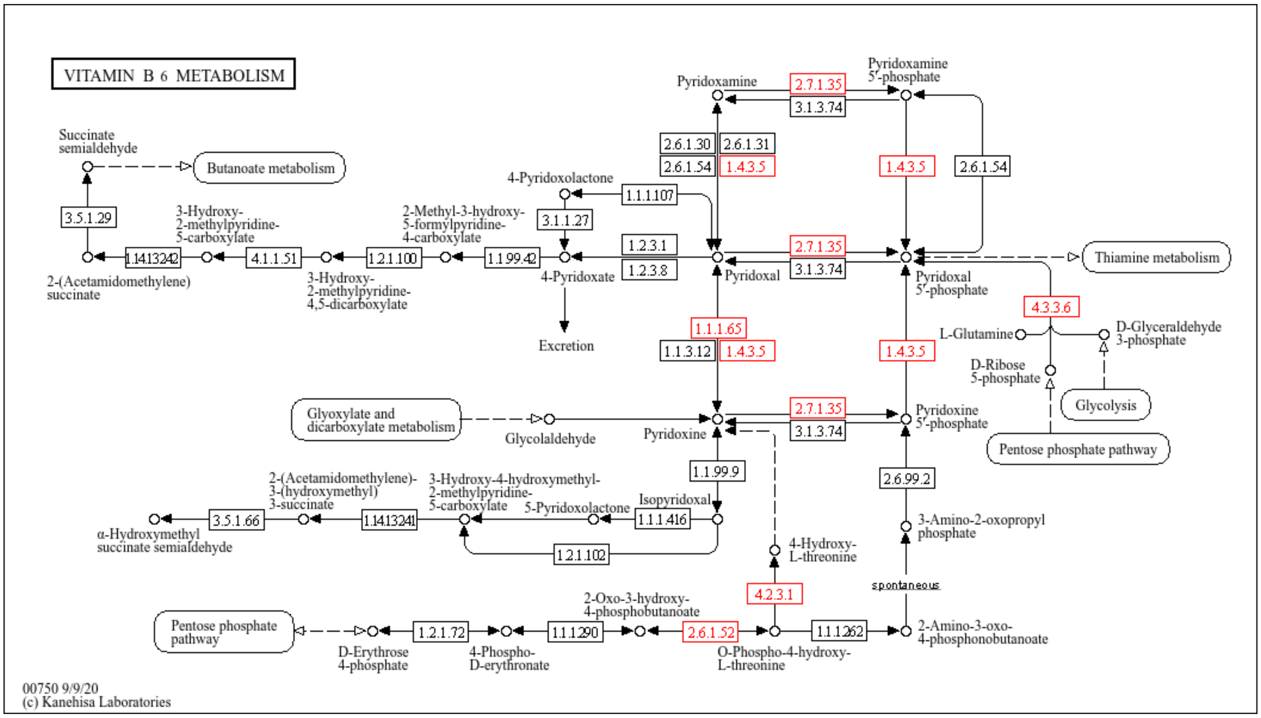

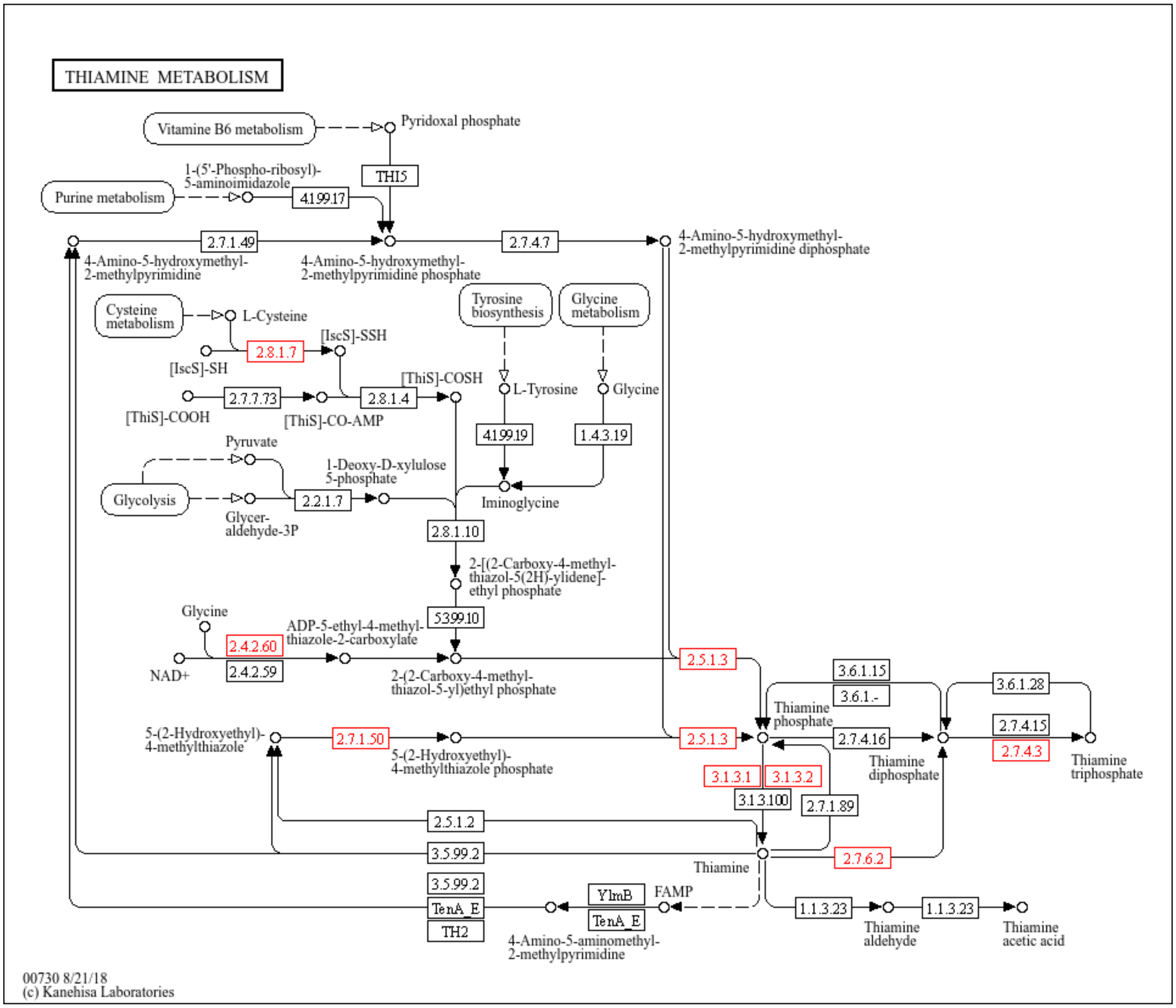
THIAMINE METABOLISM
Vitamine B6 metabolism
Pyridoxal phosphate
THI5
Purine metabolism
1-(5'-Phospho-ribosyl)-5-aminoimidazole
4.1.99.17
2.7.1.49
2.7.4.7
4-Amino-5-hydroxymethyl-2-methylpyrimidine
4-Amino-5-hydroxymethyl-2-methylpyrimidine phosphate
4-Amino-5-hydroxymethyl-2-methylpyrimidine diphosphate
Cysteine metabolism
L-Cysteine
2.8.1.7
[IscS]-SSH
[IscS]-SH
Tyrosine biosynthesis
Glycine metabolism
[ThiS]-COSH
2.7.7.73
2.8.1.4
L-Tyrosine
Glycine
[ThiS]-COOH
[ThiS]-CO-AMP
4.1.99.19
1.4.3.19
Pyruvate
1-Deoxy-D-xylulose 5-phosphate
Glycolysis
2.2.1.7
Iminoglycine
Glycer-aldehyde-3P
2.8.1.10
2-[(2-Carboxy-4-methyl-thiazol-5(2H)-ylidene]-ethyl phosphate
5.3.99.10
Glycine
ADP-5-ethyl-4-methyl-thiazole-2-carboxylate
2.4.2.60
2.4.2.59
NAD+
2-(2-Carboxy-4-methyl-thiazol-5-yl)ethyl phosphate
2.5.1.3
3.6.1.15
3.6.1.-
3.6.1.28
Thiamine phosphate
2.7.4.15
2.7.4.3
5-(2-Hydroxyethyl)-4-methylthiazole
2.7.1.50
5-(2-Hydroxyethyl)-4-methylthiazole phosphate
2.5.1.3
2.7.4.16
Thiamine diphosphate
Thiamine triphosphate
3.1.3.1
3.1.3.2
3.1.3.100
2.7.1.89
2.5.1.2
3.5.99.2
2.7.6.2
Thiamine
3.5.99.2
YlmB
FAMP
TenA_E
TenA_E
TH2
1.1.3.23
1.1.3.23
4-Amino-5-aminomethyl-2-methylpyrimidine
Thiamine aldehyde
Thiamine acetic acid
00730 8/21/18
(c) Kanehisa Laboratories

**Supplementary Figure 4**: Genomic landscapes of *Geosmithia* species.

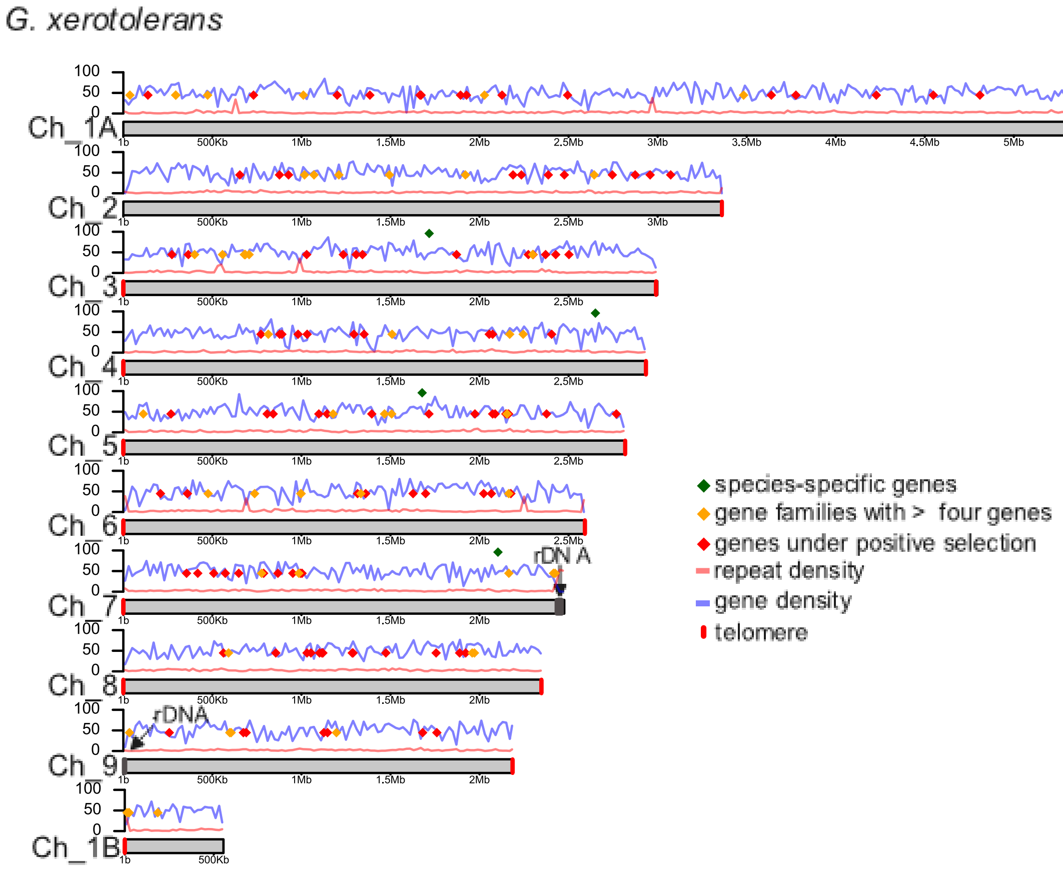

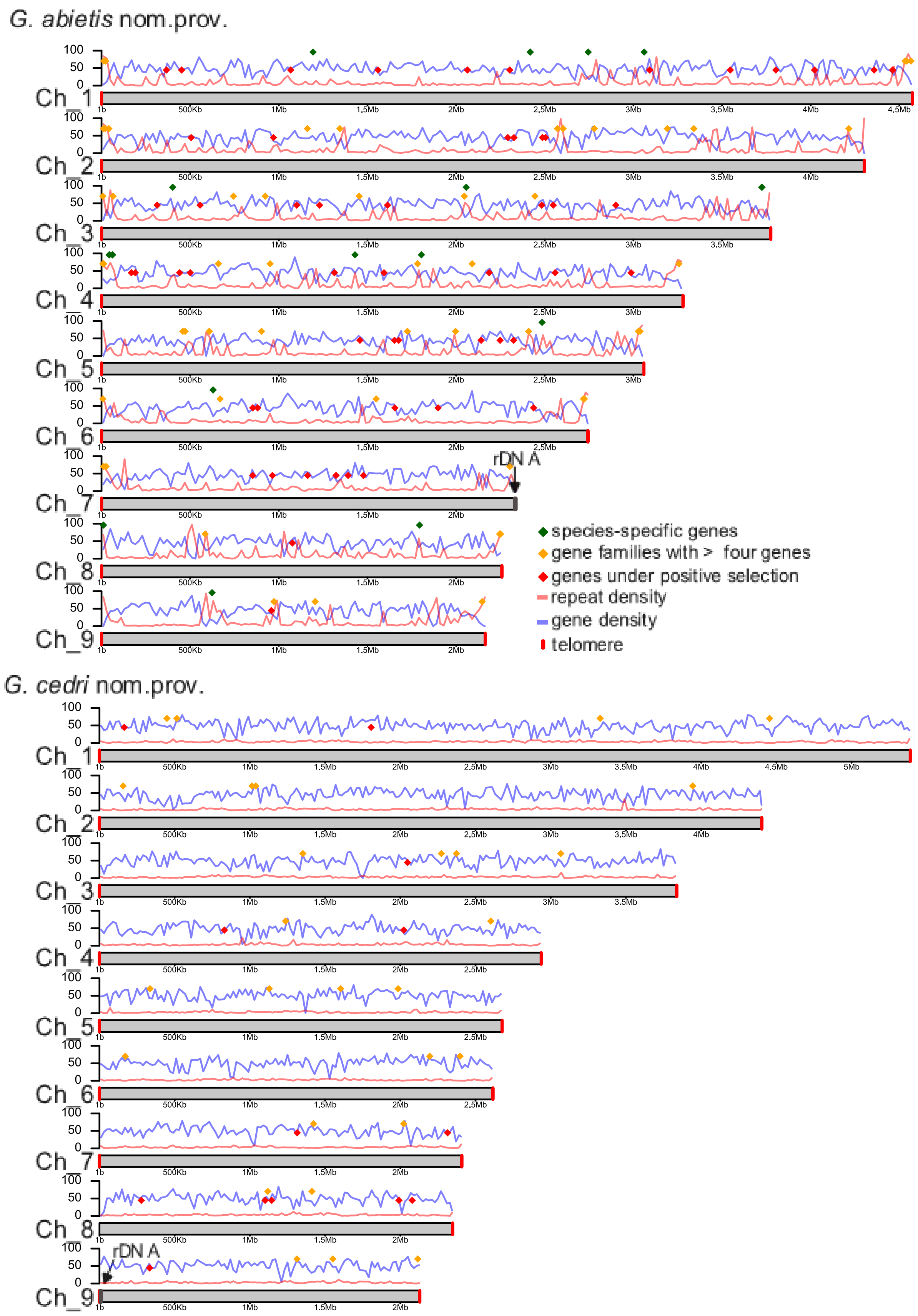

G. abietis nom.prov.
G. cedri nom.prov.
Ch_1
Ch_2
Ch_3
Ch_4
Ch_5
Ch_6
Ch_7
Ch_8
Ch_9
rDN A
species-specific genes
gene families with > four genes
genes under positive selection
repeat density
gene density
telomere

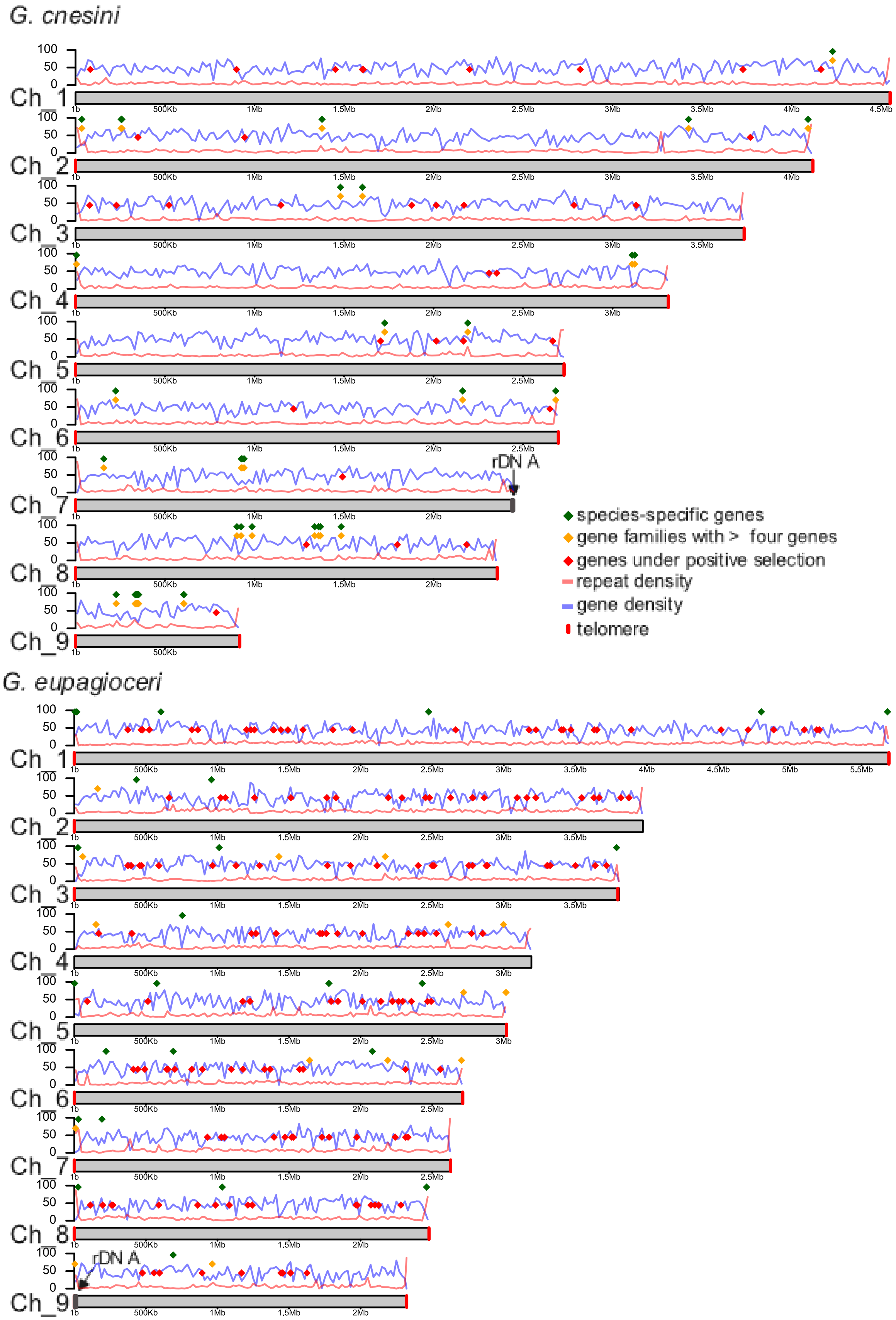
G. cnesini
Ch_1
Ch_2
Ch_3
Ch_4
Ch_5
Ch_6
Ch_7
Ch_8
Ch_9
rDN A
species-specific genes
gene families with > four genes
genes under positive selection
repeat density
gene density
telomere
G. eupagioceri
Ch_1
Ch_2
Ch_3
Ch_4
Ch_5
Ch_6
Ch_7
Ch_8
Ch_9
rDN A

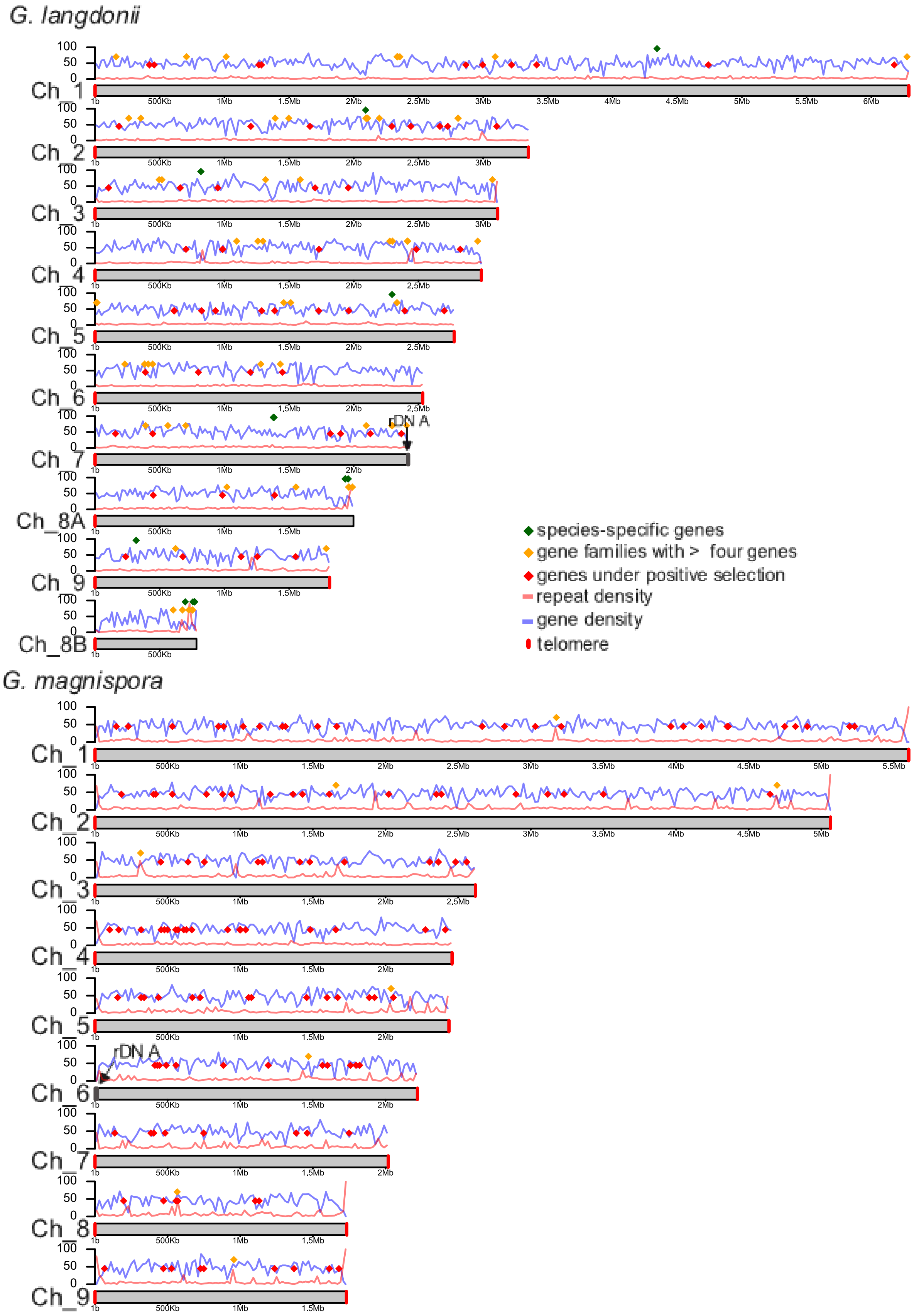

G. langdonii
Ch_1
Ch_2
Ch_3
Ch_4
Ch_5
Ch_6
Ch_7
Ch_8A
Ch_9
Ch_8B
rDNA
species-specific genes
gene families with > four genes
genes under positive selection
repeat density
gene density
telomere
G. magnispora
Ch_1
Ch_2
Ch_3
Ch_4
Ch_5
Ch_6
Ch_7
Ch_8
Ch_9
rDNA

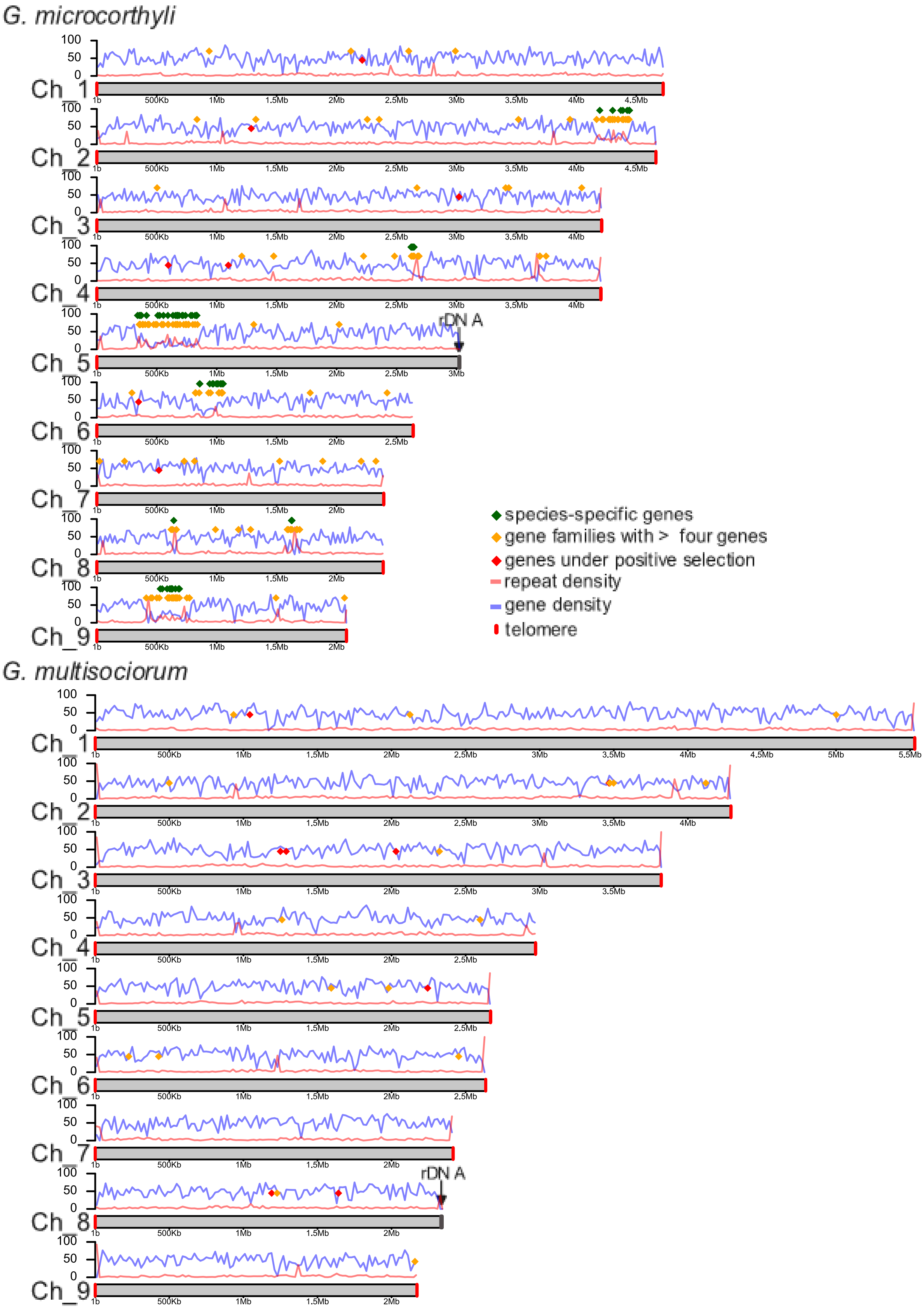

G. microcorthyli
Ch_1
Ch_2
Ch_3
Ch_4
Ch_5
Ch_6
Ch_7
Ch_8
Ch_9
rDN A
species-specific genes
gene families with > four genes
genes under positive selection
repeat density
gene density
telomere
G. multisociorum
Ch_1
Ch_2
Ch_3
Ch_4
Ch_5
Ch_6
Ch_7
Ch_8
Ch_9
rDN A

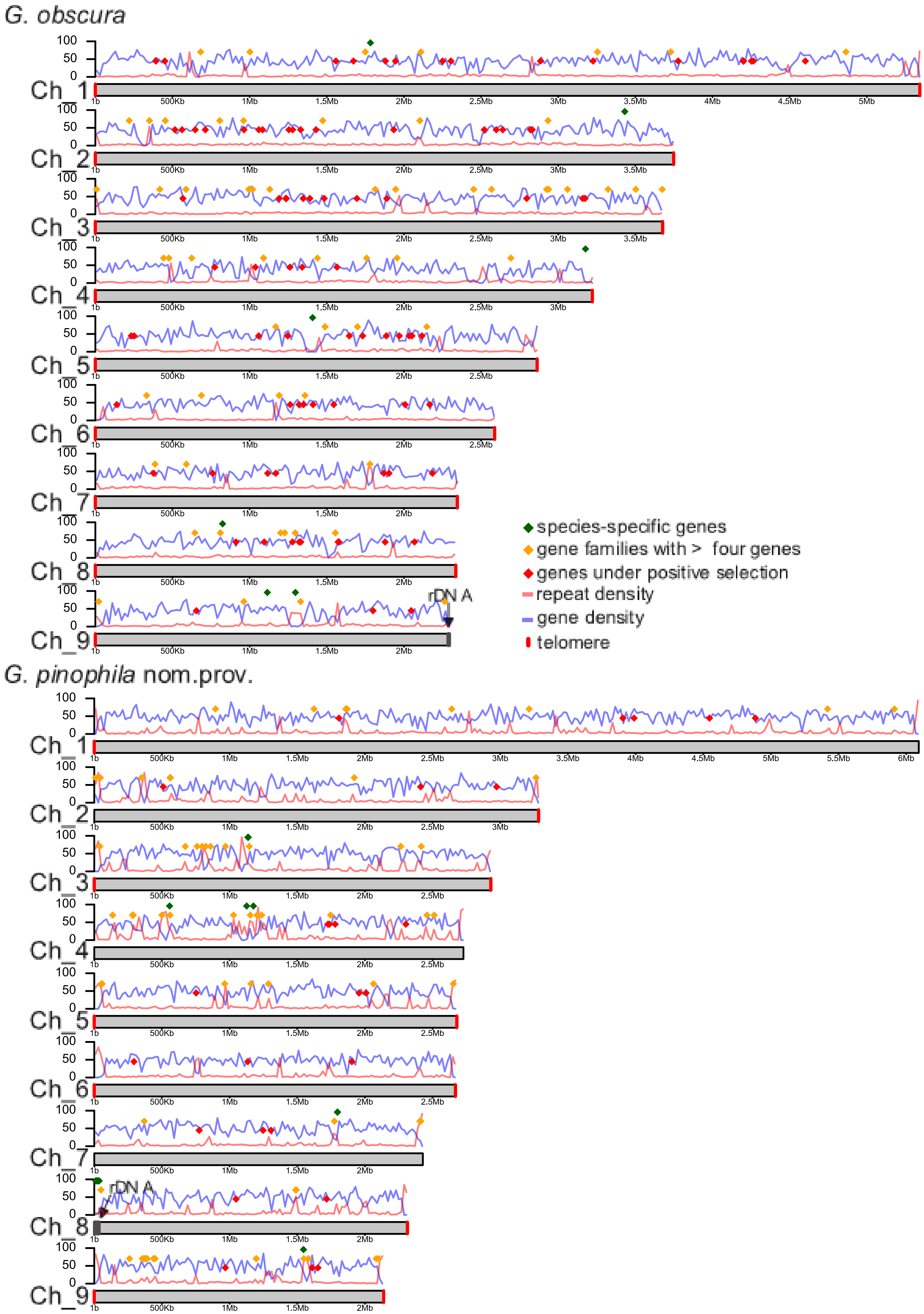
G. obscura
G. pinophila nom.prov.
Ch_1
Ch_2
Ch_3
Ch_4
Ch_5
Ch_6
Ch_7
Ch_8
Ch_9
100
50
0
1b
500Kb
1Mb
1.5Mb
2Mb
2.5Mb
3Mb
3.5Mb
4Mb
4.5Mb
5Mb
5.5Mb
6Mb
rDN A
species-specific genes
gene families with > four genes
genes under positive selection
repeat density
gene density
telomere

**Supplementary Figure 5**: Hi-C contact maps (A, C, E, G) for four *Geosmithia* species showing Rabl chromosome conformation. Blue squares denote chromosomes, black squares centromeric contacts, green squares telomeric contacts. Genomic profiles along chromosomes showing high gene expression in centromeric regions (B, D, F, H).

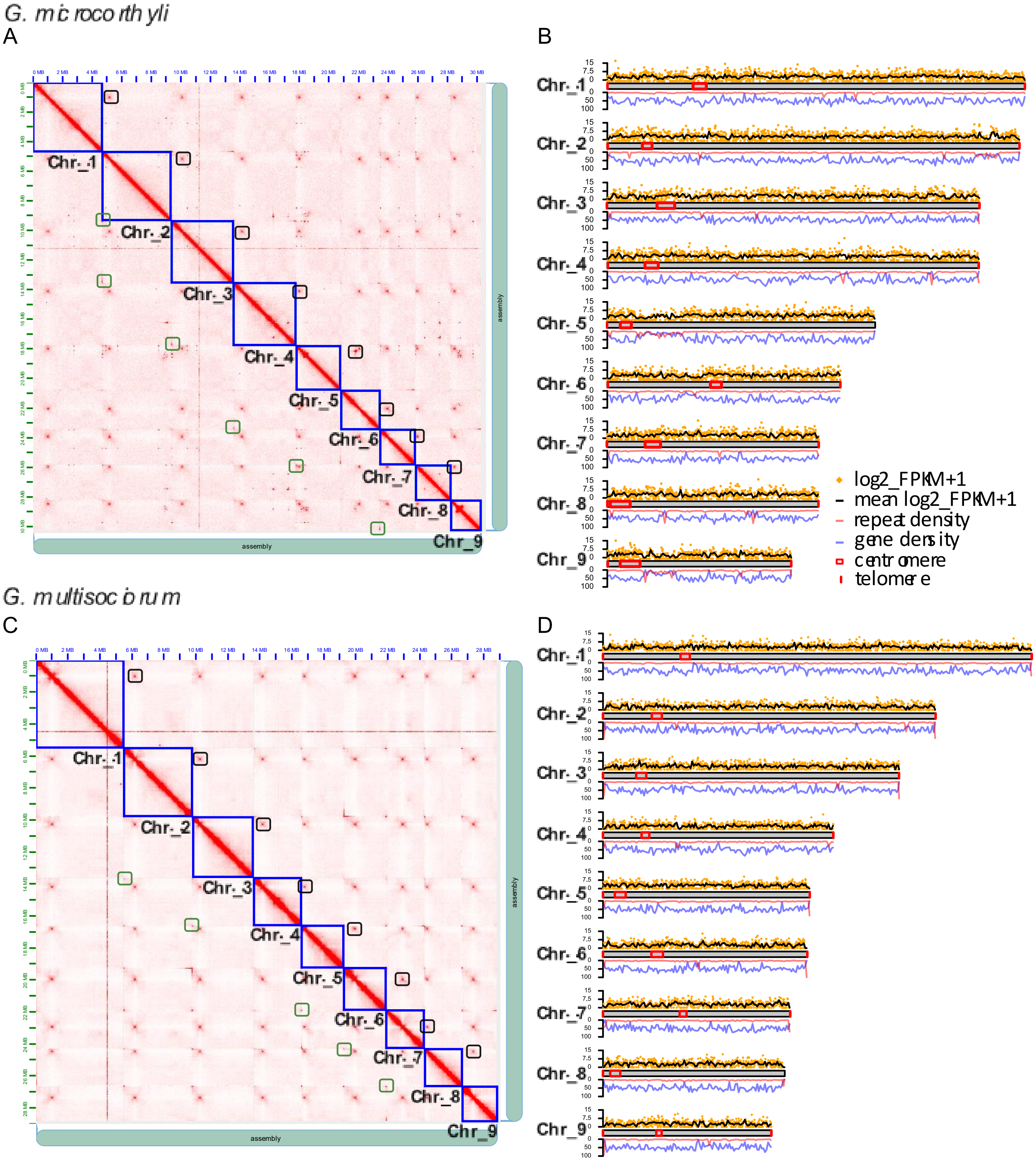

*G. eupagioceri*

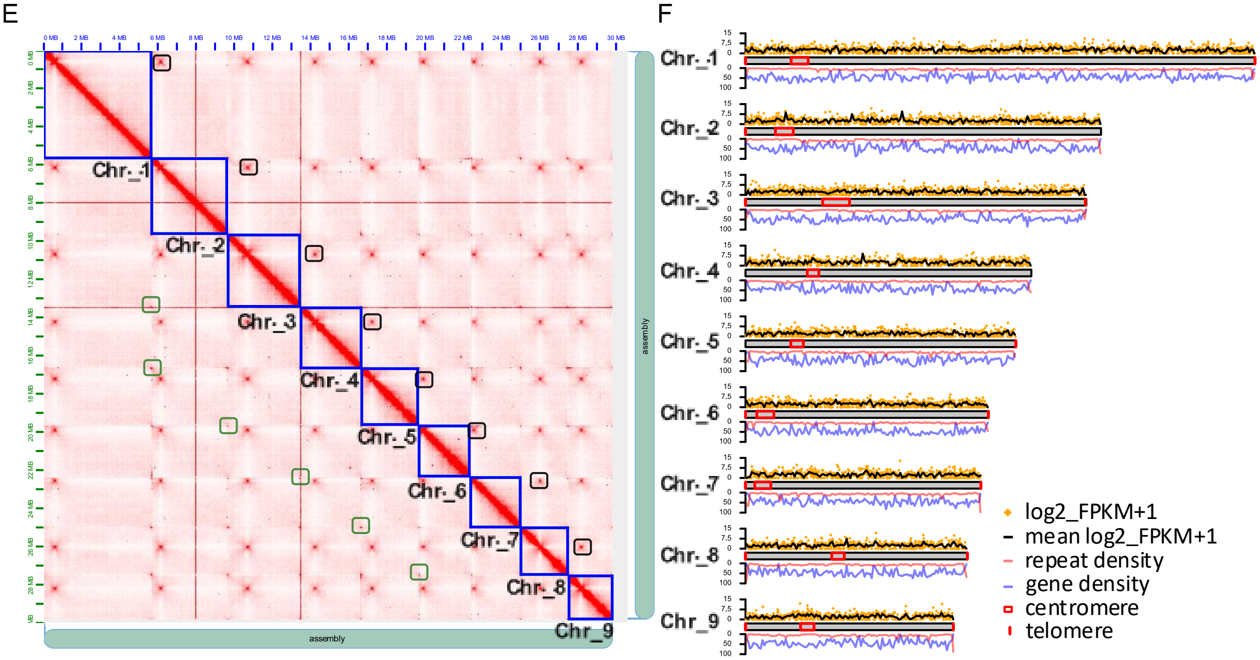


*G. pinophila nom. prov.*

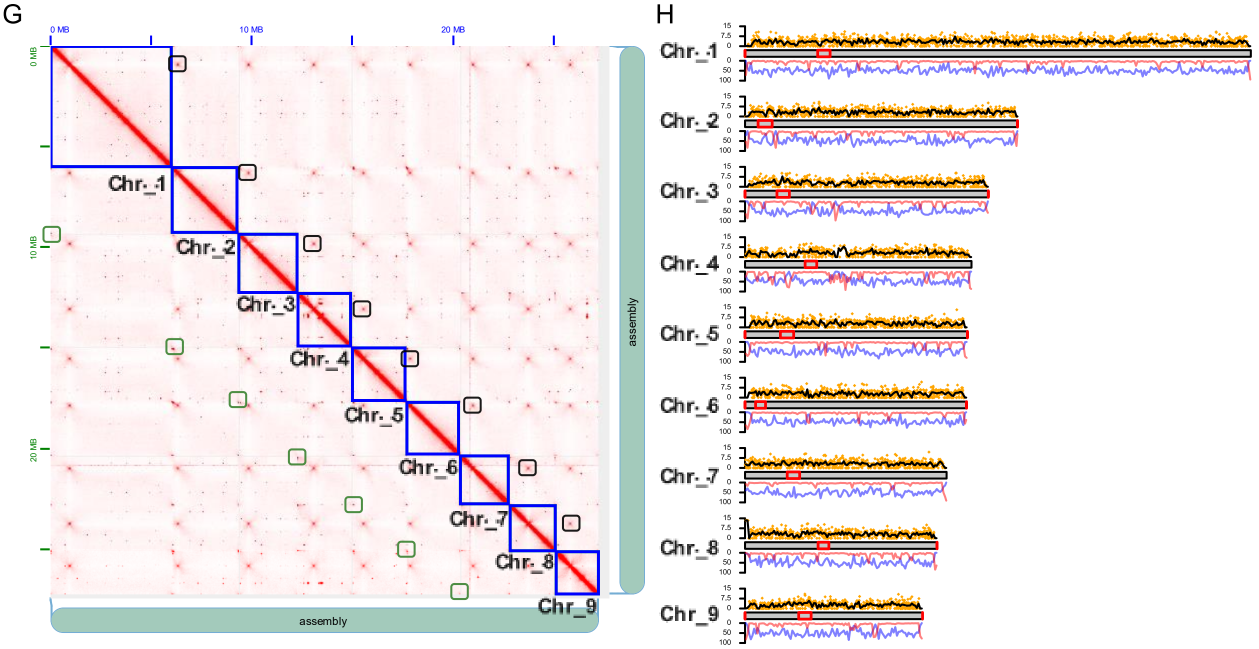

**Supplementary Table 1**. CAZyme families.

| Ecology | Species | AA | CBM | CE | GH | GT | PL |
|---|---|---|---|---|---|---|---|
| GEN | *G. cedri* nom. prov. | 62 | 9 | 9 | 165 | 81 | 7 |
| | *G. langdonii* | 61 | 9 | 10 | 165 | 82 | 7 |
| | *G. multisociorum* | 57 | 8 | 8 | 156 | 79 | 7 |
| | *G. obscura* | 55 | 7 | 11 | 155 | 79 | 6 |
| | *G. xerotolerns* | 63 | 9 | 10 | 164 | 82 | 7 |
| AMB | *G. cnesini* | 42 | 7 | 8 | 142 | 75 | 6 |
| | *G. eupagioceri* | 48 | 6 | 11 | 146 | 78 | 8 |
| | *G. microcorthyli* | 67 | 10 | 10 | 159 | 81 | 7 |
| SPE | *G. abietis* nom. prov. | 40 | 5 | 10 | 136 | 71 | 6 |
| | *G. magnispora* | 41 | 6 | 8 | 116 | 70 | 6 |
| | *G. pinophila* nom. prov. | 43 | 7 | 13 | 134 | 71 | 7 |

AA – Auxiliary Activities, CMB –Carbohydrate-Binding Modules, CE – Carbohydrate Esterases, GH – Glycoside Hydrolases, GT – Glycosyl Transferases, PL – Polysaccharide Lyases

**Supplementary Table 2**. Signal protein analysis.

| Ecology | Species | Membrane-associated | Extracellular-non effector | Effector |
|---|---|---|---|---|
| GEN | *G. cedri* nom. prov. | 204 | 295 | 108 |
| | *G. langdonii* | 203 | 301 | 103 |
| | *G. multisociorum* | 201 | 289 | 104 |
| | *G. obscura* | 207 | 286 | 96 |
| | *G. xerotolerans* | 208 | 312 | 112 |
| AMB | *G. cnesini* | 186 | 227 | 93 |
| | *G. eupagioceri* | 197 | 259 | 86 |
| | *G. microcorthyli* | 200 | 309 | 112 |
| SPE | *G. abietis* nom. prov. | 192 | 234 | 80 |
| | *G. magnispora* | 171 | 204 | 73 |
| | *G. pinophila* nom. prov. | 191 | 229 | 81 |

**Supplementary Table 3.** Secondary metabolite clusters.

| Ecology | Species | Terpene | T1PKS | NRPS-like | Indole | Fungal-RiPP-like | Hybrid |
|---|---|---|---|---|---|---|---|
| GEN | *G. cedri* nom. prov. | 5 | 5 | 7 | 0 | 0 | 2 |
| | *G. langdonii* | 5 | 6 | 5 | 1 | 1 | 1 |
| | *G. multisociorum* | 5 | 4 | 7 | 0 | 2 | 1 |
| | *G. obscura* | 5 | 5 | 5 | 1 | 1 | 2 |
| | *G. xerotolerans* | 5 | 5 | 6 | 1 | 0 | 1 |
| AMB | *G. cnesini* | 5 | 2 | 7 | 0 | 0 | 0 |
| | *G. eupagioceri* | 5 | 2 | 7 | 1 | 0 | 0 |
| | *G. microcorthyli* | 7 | 5 | 6 | 0 | 2 | 1 |
| SPE | *G. abietis* nom. prov. | 4 | 2 | 9 | 0 | 1 | 1 |
| | *G. magnispora* | 6 | 1 | 8 | 0 | 1 | 1 |
| | *G. pinophila* nom. prov. | 4 | 3 | 7 | 0 | 1 | 2 |

**Supplementary Table 4.** Pathogen-host interaction analysis.

| Ecology | Species | C | R | E | L | I | L | U | Ch |
|---|---|---|---|---|---|---|---|---|---|
| GEN | *G. cedri* nom. prov. | 2,314 | 536 | 18 | 51 | 35 | 66 | 315 | 0 |
| | *G. langdonii* | 2,254 | 921 | 23 | 50 | 40 | 75 | 442 | 1 |
| | *G. multisociorum* | 2,256 | 905 | 20 | 49 | 39 | 72 | 440 | 1 |
| | *G. obscura* | 2,205 | 874 | 20 | 50 | 40 | 72 | 448 | 1 |
| | *G. xerotolerans* | 2,269 | 904 | 21 | 49 | 38 | 75 | 442 | 1 |
| AMB | *G. cnesini* | 1,995 | 816 | 18 | 48 | 39 | 73 | 406 | 1 |
| | *G. eupagioceri* | 2,046 | 853 | 21 | 49 | 35 | 70 | 427 | 1 |
| | *G. microcorthyli* | 2,332 | 922 | 20 | 49 | 38 | 74 | 459 | 1 |
| SPE | *G. abietis* nom. prov. | 1,981 | 821 | 19 | 53 | 39 | 70 | 426 | 1 |
| | *G. magnispora* | 1847 | 798 | 16 | 48 | 29 | 73 | 388 | 1 |
| | *G. pinophila* nom. prov. | 2,028 | 827 | 19 | 49 | 42 | 73 | 409 | 1 |

C – conflict, R – reduced, E – effector, L – lethal, I – increased, L – loss, U – unaffected, Ch – chemistry.

**Supplementary Table 5**. The output from OrthoFinder analysis.

| Ecology | Species | Genes in OGs [%] | OGs containing species [%] | Number of species-specific OGs | Number of genes in species-specific OG | Single copy OG [%] | Multi copy OG [%] |
|---|---|---|---|---|---|---|---|
| GEN | *G. cedri* nom. prov. | 99.6 | 92 | 0 | 0 | 92.3 | 7.3 |
| | *G. langdoniii* | 99.1 | 89.8 | 9 | 18 | 90.5 | 8.7 |
| | *G. multisociorum* | 99.7 | 90.8 | 0 | 0 | 93.1 | 6.6 |
| | *G. obscura* | 99 | 88.6 | 3 | 7 | 91 | 7.8 |
| | *G. xerotolerans* | 99.2 | 90.7 | 2 | 4 | 91.2 | 8.1 |
| AMB | *G. cnesini* | 99.2 | 82.7 | 5 | 11 | 94.4 | 4.9 |
| | *G. eupagioceri* | 98.7 | 84.8 | 9 | 19 | 92.4 | 6.2 |
| | *G. microcorthyli* | 99.4 | 92 | 20 | 65 | 88.9 | 10.4 |
| SPE | *G. abietis* nom. prov. | 98.8 | 84.3 | 9 | 21 | 93.5 | 5.3 |
| | *G. magnispora* | 98.9 | 78.4 | 0 | 0 | 94.5 | 4.5 |
| | *G. pinophila* nom. prov. | 99 | 84.1 | 5 | 12 | 93.3 | 5.6 |
